# Identifying Novel Roles for Peptidergic Signaling in Mice

**DOI:** 10.1101/675603

**Authors:** Kathryn G. Powers, Xin-Ming Ma, Betty A. Eipper, Richard E. Mains

## Abstract

Despite accumulating evidence demonstrating the essential roles played by neuropeptides, it has proven challenging to use this information to develop therapeutic strategies. Peptidergic signaling can involve juxtacrine, paracrine, endocrine and neuronal signaling, making it difficult to define physiologically important pathways. One of the final steps in the biosynthesis of many neuropeptides requires a single enzyme, peptidylglycine α-amidating monooxygenase (PAM), and lack of amidation renders most of these peptides biologically inert. PAM, an ancient integral membrane enzyme that traverses the biosynthetic and endocytic pathways, also affects cytoskeletal organization and gene expression. While mice, zebrafish and flies lacking *Pam* (*Pam*^KO/KO^) are not viable, we reasoned that cell-type specific elimination of *Pam* expression would generate mice that could be screened for physiologically important and tissue-specific deficits. *Pam*^cKO/cKO^ mice, with loxP sites flanking the 2 exons deleted in the global *Pam*^KO/KO^ mouse, were indistinguishable from wildtype mice. Eliminating *Pam* expression in excitatory forebrain neurons reduced anxiety-like behavior, increased locomotor responsiveness to cocaine and improved thermoregulation in the cold. A number of amidated peptides play essential roles in each of these behaviors. Although atrial natriuretic peptide (ANP) is not amidated, *Pam* expression in the atrium exceeds levels in any other tissue. Eliminating *Pam* expression in cardiomyocytes increased anxiety-like behavior and improved thermoregulation. Atrial and serum levels of ANP fell sharply *Pam*^Myh6-cKO/cKO^ in mice and RNASeq analysis identified changes in gene expression in pathways related to cardiac function. Use of this screening platform should facilitate the development of new therapeutic approaches targeted to peptidergic pathways.

**SIGNIFICANCE:** Peptidergic signaling, which plays key roles in the many pathways that control thermoregulation, salt and water balance, metabolism, anxiety, pain perception and sexual reproduction, is essential for the maintenance of homeostasis. Despite the fact that peptides generally signal through G protein coupled receptors, it has proven difficult to use knowledge about peptide synthesis, storage and secretion to develop effective therapeutics. Our goal was to develop an *in vivo* bioassay system that would reveal physiologically meaningful deficits associated with disturbed peptidergic signaling. We did so by developing a system in which an enzyme essential for the production of many bioactive peptides could be eliminated in a tissue-specific manner.

## INTRODUCTION

Peptidergic signaling plays a key role in the vertebrate nervous system and in animals such as *Trichoplax adhaerens*, which lack neurons and muscles (1). Transcripts encoding putative *T. adhaerens* preproneuropeptides are expressed in a cell-type specific manner, and peptides that could be derived from them govern movement and contractile behavior (2). Acting as co-transmitters and neuromodulators in organisms with a nervous system, neuropeptides play critical roles in the complex pathways that control pain perception, mating behaviors, anxiety, appetite and metabolism (3).

Bioactive neuropeptide synthesis in *Trichoplax* and mammals involves a similar set of post-translational modifications, each of which occurs in the lumen of the secretory pathway (4–7). The bioactive peptides are stored in secretory granules and released in response to an appropriate stimulus. The vast majority of neuropeptides act through G protein coupled receptors (GPCRs), which may be located close to or far from the site at which the bioactive peptide is released (8). Neuropeptides were identified using bioassays, biochemical paradigms, screens for GPCR ligands and genetic selections. Some bioactive peptides, such as neuropeptide Y (NPY), were identified by the presence of a common post-translational modification, C-terminal amidation, which is often essential for bioactivity (8, 9).

Preproneuropeptides that yield amidated products encode precursors that generate a peptidylglycine intermediate. Peptidylglycine α-amidating monooxygenase (PAM; E.C. 1.4.17.3) catalyzes the two step conversion of the penultimate residue of its peptidylglycine substrate into an α-amide, releasing glyoxylate (8). Cell culture experiments have revealed additional roles for PAM in regulating cytoskeletal organization, secretagogue responsiveness and gene expression (10). PAM knockout mice (*Pam*^KO/KO^), zebrafish and *Drosophila* are not viable (11–13). *Pam*^KO/KO^ mouse embryos develop pericardial edema, have a poorly developed yolk sac vasculature and do not survive beyond the second week of gestation (11). Mice with a single copy of *Pam* are sensitive to seizures, exhibit increased anxiety-like behavior and cannot maintain body temperature in the cold (14). Genetic studies indicate that *PAM* is associated with diabetes, Alzheimer disease, Parkinson disease, hypertension and pituitary tumors (reviewed in (8)).

We hypothesized that generation of a mouse line in which *Pam* expression can be eliminated in a cell type-specific manner would facilitate the identification of behaviors and metabolic pathways in which neuropeptides and PAM play significant roles. LoxP sites were placed around the exons eliminated in the *Pam*^KO/KO^ mouse (11), generating the *Pam*^cKO/cKO^ mouse. Knowing that behaviors rely on precisely balanced inputs from excitatory and inhibitory neurons, both of which express PAM (15), we bred *Pam*^cKO/cKO^ mice to mice expressing Cre-recombinase in their forebrain excitatory neurons, under control of the spiracles homeobox 1 promoter (Emx-1 Cre) (16, 17). Given the high levels of PAM expressed in atrial cardiomyocytes (18), we also bred *Pam*^cKO/cKO^ mice to mice expressing Cre-recombinase under control of the cardiac myosin heavy chain 6 promoter (Myh6-Cre) (19, 20). Tests of anxiety-like behavior and thermal regulation demonstrated alterations in both lines. Although atrial natriuretic peptide (ANP) is not amidated, ANP levels fell dramatically in *Pam*^*Myh6-cKO/cKO*^ mice and gene expression was altered.

Using the *Pam*^cKO/cKO^ mouse, PAM expression and amidated peptide levels can be altered in selected cell types when developmentally or experimentally expedient. In this way, it should be possible to identify the circuits and complex peptidergic pathways that play key roles behaviors of interest.

## METHODS

### Construction of the *Pam*^cKO/cKO^ mouse

The *Pam*^cKO/cKO^ mouse was created directly on a C57BL6 background by Cyagen Biosciences (Santa Clara, CA), closely following the original *Pam* global (whole animal, not tissue- or cell-specific) knockout targeting Exons 2 and 3 on mouse Chromosome 1 (*Pam*^KO/KO^) (11). The upstream loxP site is 300 nt before the start of exon 2, bracketed by the *Pam* forward and reverse primers (**Supplemental Table 1**), and the downstream loxP site is 600 nt beyond the end of Exon 3 (**Supplemental Fig.1A**). The *Pam* deletion primer is 2.7 kb from the *Pam* forward primer on the mouse genome, so the intervening DNA has to be removed for the outside primers to make the 310 nt product in the 1 min polymerase chain reaction (PCR) elongation step. The three primer reaction makes possible genotyping distinguishing wildtype (WT), *Pam*^cKO/+^ and *Pam*^cKO/cKO^ mice in one reaction, using DNA from identification earpunches. Tissue-specific verification of the excision of Exons 2+3 was also demonstrated with the same primers and PCR program: 94C, 3m; 94C, 30s; 60C, 35s; 72C, 1m; 32 cycles. The *Pam*^cKO/+^ and *Pam*^cKO/cKO^ mice were born in the expected Mendelian ratios, were fertile and exhibited no obvious abnormalities (see below).

### Mice expressing Cre-recombinase

Two lines of Cre-recombinase mice were purchased from Jackson Laboratories (Bar Harbor, ME). Excitatory cortical neuron-specific excision was targeted using JAX005628-B6.129S2-*Emx1*^*tm1(cre)Krj*^/J, maintained on a C57BL/6J background as homozygotes (16). Convenient genotyping required sequencing the 3’UTR of the Emx1 gene in the Emx1-Cre mice, to make possible three-primer screening (**Supplemental Fig.1B; Suppl.Table 1**). Three primer reactions distinguished WT, heterozygous and homozygous Emx1-Cre mice (94C, 3m; 94C, 30s; 61C, 30s; 72C,1m; 30 cycles). As noted on the JAX website, germline expression of Cre recombinase occurred at a low but significant frequency in mice expressing Emx1-Cre and one floxed PAM allele. Evidence of this conversion was observed in ear punches and tail clips as well as verified with multiple brain and heart tissues genotyped using the *Pam*^cKO^ primer set. The mice with *Pam* recombination of a single allele in all tissues were similar to the previously studied *Pam*^KO/+^ global heterozygote knockout mice (11, 14, 21) in every tested biochemical, physiological and behavioral parameter. These *Pam*^KO/+^ mice were thus excluded from the current analyses. Excitatory neuron-specific PAM knockout mice were produced with Emx1-Cre expressed in one or both parents, since there was no evidence of any problems with Emx1-Cre homozygosity. Excitatory neuron-specific *Pam* knockouts are designated *Pam*^Emx1-cKO/cKO^ and heterozygotes are *Pam*^Emx1-cKO/+^.

Cardiomyocyte-specific expression of Cre-recombinase was achieved using JAX011038-B6.FVB-Tg(Myh6-cre)2182Mds/J, maintained on a C57BL/6J background as hemizygotes (19). Genotyping was performed with a primer set closely modeled on the primers suggested on the JAX website (**Supplemental Fig.1C; Suppl.Table 1**); the primers were modified to enable the use of three primers in a single definitive genotyping for each animal (94C, 3m; 94C, 30s; 62C, 30s; 72C,45s; 30 cycles). As suggested on the JAX website, cardiomyocyte-specific *Pam* knockout mice (*Pam*^Myh6-cKO/cKO^) were produced with Myh6-Cre expressed in only one parent, to avoid the potential toxicity of Myh6-Cre homozygosity. Cardiac-specific *Pam* knockouts are designated *Pam*^Myh6-cKO/cKO^ and heterozygotes are *Pam*^Myh6-cKO/+^. Heart weight was unaltered in adult *Pam*^Myh6-cKO/cKO^ mice.

### Biochemical analyses

Lysates for PHM and PAL assays were prepared by homogenizing tissue in 20 mM Na TES, 10 mM mannitol, 1% TX-100, pH 7.4 (TMT), with protease inhibitors; all manipulations were carried out on ice, as described (22). Lysates for SDS-PAGE were prepared using SDS-P buffer with protease inhibitor cocktail; after boiling and sonication, particulate material was removed (23). Lysates were also prepared by sonication into ice cold RIPA Buffer (Cell Signaling Technologies, Danvers, MA; #9806) containing protease inhibitors (Sigma P8340, MilliporeSigma, Burlington MA); particulate material was then removed. Protein concentrations were determined using the bicinchoninic acid assay (Thermo Fisher, Waltham MA), with bovine serum albumin as the standard. For PHM and PAL activity assays, samples were diluted to 0.1 mg/ml and assayed in triplicate. Amounts and assay times were adjusted to ensure assay linearity: atrium (0.2 μg, 30 min), ventricle (1.0 μg, 120 min), cortex (5.0 μg, 60 min), hypothalamus (2.0 μg, 60 min), olfactory bulb (1.0 μg, 60 min).

For Western blots, samples were denatured in SDS sample buffer by heating at 95°C for 5 min: for atrium, 5 or 10 μg of protein was loaded; for all other tissues, 10 or 20 μg of protein was loaded (22). Samples fractionated on Bio-Rad Criterion TGX Precast 4-15% gradient gels were transferred to Immobilon-P transfer membranes (Millipore). Membranes were blocked with 5% milk in TTBS before overnight incubation with primary antibody and visualization using HRP-tagged secondary antibody (Jackson ImmunoResearch Laboratories, West Grove, PA) and SuperSignal West Pico PLUS Chemiluminescent substrate (ThermoFisher). ProANP was detected using rabbit polyclonal antibody to proANP(1-16) (kindly provided by Dr. Christopher Glembotski, San Diego State University) (24, 25); ANP was detected with a goat antibody (ab190001, Abcam, Cambridge MA). PAM was visualized using affinity-purified rabbit polyclonal antibodies to the linker region separating PHM from PAL in PAM-1 (JH629; RRID:AB_2721274) (22), and to the extreme cytoplasmic domain of PAM (C-Stop: RRID:AB_2801640) (26). In addition to using Coomassie Brilliant Blue staining to verify equal loading, Iqgap was visualized using a mouse monoclonal antibody (#610611, BD Transduction Laboratories, San Jose, CA). Western blot signals in the linear range were densitized using GeneTools software (Syngene, Frederick MD).

The ELISA for ANP was performed as described (EIAM-ANP-1, RayBiotech, Peachtree Corners, GA), except that the standard curve was modified to include 4-fold dilutions from 80 pg to 0.0195 pg; values were calculated using a logit-log transformation. Tissue and serum samples were prepared as described by de Bold, Smithies and coworkers (27, 28) using UltraMicroSpin C18 silica columns (Nest Group, Southborough, MA). Serum renin levels were determined using an ELISA kit (ELM-Renin1-1, RayBiotech).

### Tissue and cellular specificity of PAM ablation

As an initial determination of the sites of PAM depletion, Myh6-Cre and Emx1-Cre mice were individually crossed with TdTomato reporter mice, and *Pam*^Emx1-cKO/cKO^ mice were studied directly (29). Direct observation of PAM at the cellular level was accomplished by immunocytochemistry as described (30) using affinity-purified PAM polyclonal antibody JH629. Nuclei were visualized with Hoechst stain and, where indicated, Gad67 was visualized using a mouse monoclonal antibody (clone 1G10.2, Millipore) (31). Confocal imaging was performed on 12 μm sections with a Zeiss LSM 880 microscope with a 20x, 40x, or 63x objective; where indicated, z-stacks were taken and a single image is shown.

### Behavioral and physiological analyses

All experiments were performed in the same circadian period (0900-1600). Mice were handled by the same animal tester for 60 seconds a day for two days before determining the fraction of time in the open area of the elevated zero maze (San Diego Instruments, San Diego, CA) (32). Mice were video recorded once for 5 min; times were obtained by watching the movies. Next, the general mobility and motor coordination of the mice were tested using the open field and rotarod (29, 32). Rotarod testing was performed three times a day for 3 days, using a Five Lane Rota-Rod for Mouse (Med Associates, Georgia, VT). Each trial lasted no longer than 5 min, with the speed increasing from 4 rpm to 40 rpm. The longest time recorded for each day was used in the analysis. Open field ambulations were recorded for 15-45 min (see Figure Legends) in a Photobeam Activity System Open Field (San Diego Instruments). When the locomotor response to cocaine was tested, cocaine was administered by intraperitoneal injection and open field ambulations were recorded for 45 min on successive days or after a brief withdrawal period; cocaine injections were 10 mg/kg except for day 2, which was 20 mg/kg (33, 34).

Core body temperature was recorded over a 3-hour period using a rectal thermometer (14). Temperature readings were collected before entering the 4°C cold room and once each following hour in the cold room. During the 3h experiment, mice were singly housed in plain pre-chilled mouse cages with no bedding, food or water. Abdominal and tail temperatures during 20 min restraint in an unfamiliar environment (ventilated 50 ml tube) at room temperature were determined using a Hti Dual Laser Infrared Thermometer (35, 36). Blood pressure was determined using the tail cuff method (CODA Multi Channel, Computerized, Non-Invasive Blood Pressure System for Mice and Rats; Kent Scientific Corporation, Torrington, CT) (28, 37, 38); arterial blood pressures by tail cuff method or telemetry are identical with experienced operators (37). Dietary NaCl was varied using standard mouse chows from Envigo (Huntingdon, UK): diet TD.96208 contained 0.49% NaCl; the high salt diet TD.92012 contained 8% NaCl. The standard chow in the University of Connecticut Health Center Animal Tower was the irradiated version of the Teklad Global 18% Rodent Diet (2918) from Envigo and contained 0.49% NaCl, which we refer to as a normal salt diet. The effect of elimination of sympathetic nervous system input on systemic blood pressure was determined 10 min after intraperitoneal injection of 0.9% saline (control) or hexamethonium (30 mg/kg) (# 4111, Tocris, Minneapolis, MN) in saline (38, 39).

For behavioral testing, control animals (CON) included wildtype mice and mice with one or two floxed *Pam* alleles but no Cre-recombinase, or Cre-recombinase but no floxed *Pam* alleles, since these mice were behaviorally and biochemically indistinguishable from wildtype mice. Only wildtype mice were used as controls in physiological testing (e.g. blood pressure). Since males and females did not differ within a genotype except for body weight and blood pressure values, data were pooled across sexes for each genotype.

### RNA-Sequencing

RNA prepared from individual adult mouse atria was sequenced and aligned as described (10, 34). Sequencing was paired-end with 4.7 to 7.4 million reads mapped to mm10 per sample. Differential expression was analyzed using DESeq2 analyses (40, 41) (https://bioconductor.org/packages/release/bioc/html/DESeq2.html) (cutoff p<0.05), followed by removal of outliers (10, 34). RNASeq data were analyzed using Ingenuity Pathway Analysis (Qiagen, Germantown, MD) (10, 34). Sequencing data are available online (GEO Submission GSE132180).

### Data and Unique Materials Availability

All data, protocols and reagents (e.g. antisera) are available upon request to. The *Pam*^cKO/cKO^ mice are available on request by qualified researchers for their own use. These mice will be deposited at JAX as soon as possible.

### Statistical analyses

Student t-tests were applied in pairwise comparisons of enzyme activity or protein abundance (Excel or OpenOffice). Time courses (e.g. growth curves) were subjected to 2-way ANOVA (Prism 8). RNAseq differential expression was evaluated using DESeq2, which has its own built-in statistical comparisons.

## RESULTS

### Mice lacking *Pam* expression driven by Emx1-Cre or Myh6-Cre grow and perform locomotor tasks normally

PAM activity is detected in mouse embryos by embryonic day 12.5 (E12.5); embryos unable to express *Pam* are not found after E14.5 (11). Since Emx1-Cre is active by E10.5 (16) and an Myh6-LacZ reporter is expressed in the E9.5 atrium (20), it is likely that Cre-recombinase mediated excision of *Pam* exons 2 and 3 occurs before expression of *Pam* would have begun. In the tissues and sera of adult heterozygous global *Pam* knockout mice (*Pam*^KO/+^), PAM activity and protein levels are approximately half of wildtype values (11).

For both tissue-specific *Pam*^cKO/cKO^ lines, females grew normally from weaning to 100 days of age (**Figs.1A,B**); *Pam*^*Emx1-cKO/cKO*^ male mice did not differ from WT males, while *Pam*^*Myh6-cKO/cKO*^ males were slightly larger than WT [p<0.0001, 2-way ANOVA]. When *Pam*^*Emx1-cKO/+*^ mice were mated with *Pam*^cKO/+^ mice, the genotypes of progeny tested at weaning occurred in the expected Mendelian ratio (N=226; N.S. by 2-way ANOVA) (**Supplemental Fig.2A**). Similarly, when Myh6-Cre progeny were tested, the expected Mendelian ratio of genotypes was observed (N=198; N.S. by 2-way ANOVA) (**Supplemental Fig.2B**).

**Fig.1.**
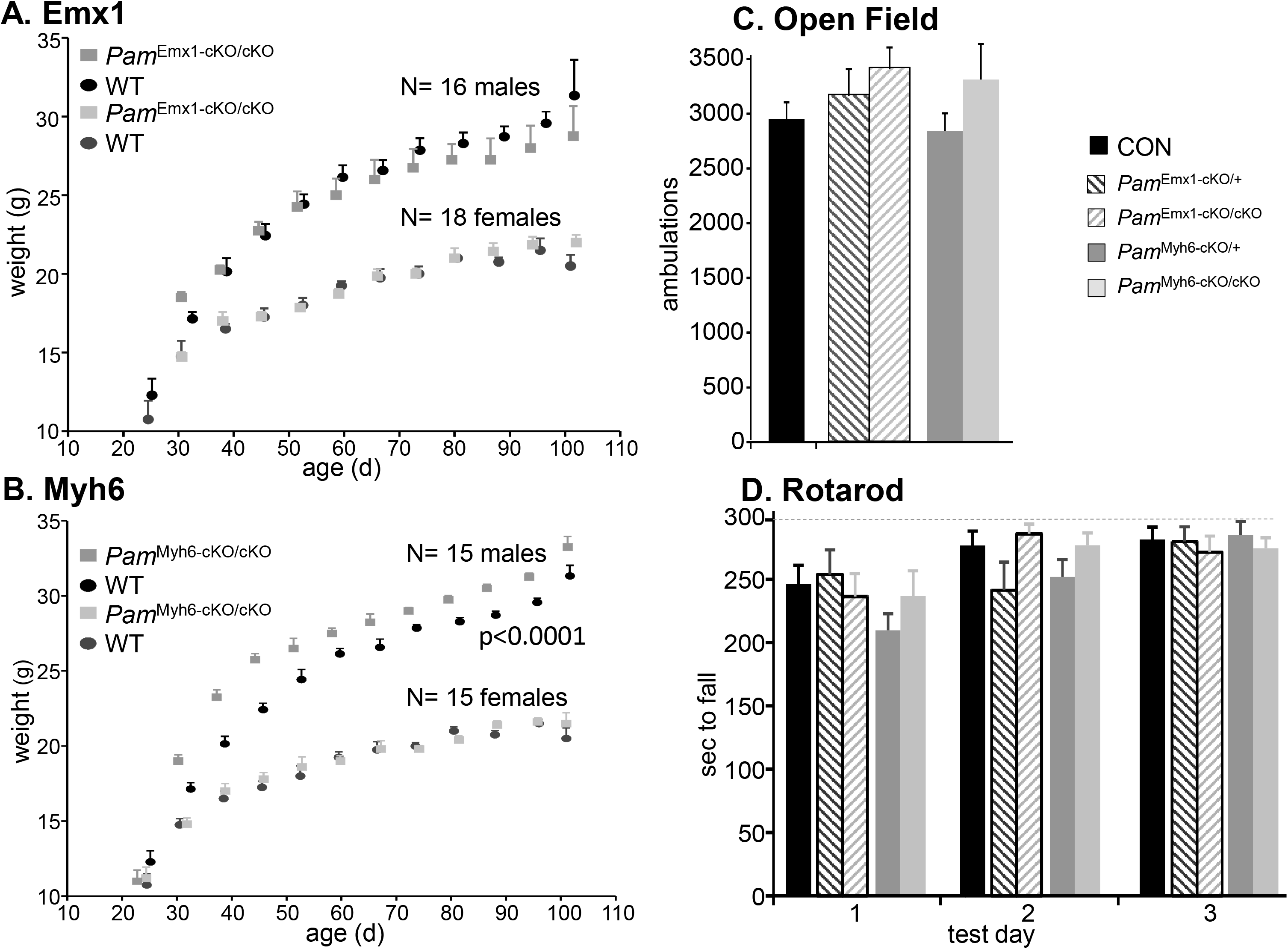
Mice from both Cre-driver lines grow and move normally. **Growth Curves. A.** The growth of male (N=15) and female (N=16) *Pam*^Emx1-cKO/cKO^ mice was indistinguishable from wildtype (WT) littermates. **B.** *Pam*^Myh6-cKO/cKO^ males were larger than wildtype male littermates (N=16), while the corresponding females were indistinguishable from wildtype females (N=18). **C. Open Field locomotor activity.** Ambulations (beam breaks) were recorded over a 30 min period: CON, n = 43; *Pam*^Emx1-cKO/+^, n = 22; *Pam*^Emx1-cKO/cKO^, n = 24; *Pam*^Myh6-cKO/+^, n = 26; *Pam*^Myh6-cKO/cKO^, n = 19. For **C** and **D**, males and females were pooled since they were indistinguishable. **D. Rotarod**. Longest time to fall from the accelerating Rotarod on each test day is plotted: Control (CON), n = 22; *Pam*^Emx1-cKO/+^, n = 8; *Pam*^Emx1-cKO/cKO^, n = 16; *Pam*^Myh6-cKO/+^, n = 16; *Pam*^Myh6-cKO/cKO^, n = 8.

Since most behavioral tests require normal mobility, we tested male and female *Pam*^Emx1-cKO/cKO^ and mice on the Rotarod and in the Open Field. Since no differences were observed in the behavior of wildtype, *Pam*^cKO/cKO^, *Pam*^cKO/+^ and Emx1-Cre or Myh6-Cre mice, all were used as controls (CON). The *Pam*^Emx1-cKO/cKO^ and *Pam*^Myh6-cKO/cKO^ mice were as active as control mice in the Open Field (**Fig.1C**); no differences were observed in the performance of male and female mice. Male and female *Pam*^Emx1-cKO/cKO^ and *Pam*^Myh6-cKO/cKO^ mice performed as well as control mice on the Rotarod (**Fig.1D**); no differences were observed between male and female mice.

### *Pam* knockout is tissue-specific and effective

We used assays for PHM and PAL enzyme activity to assess the success of our knockout strategies. In order to extract both soluble and membrane forms of PAM, tissue from adult male and female mice was homogenized in lysis buffer containing 1% TX-100; similar results were obtained using both assays (**Figs.2A,B**). PHM and PAL specific activities were reduced to 25% of wildtype values in cortical and olfactory bulb lysates prepared from *Pam*^Emx1-cKO/cKO^ mice. A less extreme reduction in PHM and PAL specific activities was also observed in hypothalamic lysates prepared from *Pam*^Emx1-cKO/cKO^ mice. PHM and PAL specific activities were reduced to 2% of wildtype values in the atria of *Pam*^Myh6-cKO/cKO^ mice. PHM and PAL specific activities, which are much lower in the adult ventricle, were reduced to 25% of wildtype values in *Pam*^Myh6-cKO/cKO^ mice. In contrast, the specific activities of both enzymes in the atria and ventricles of *Pam*^Emx1-cKO/cKO^ mice were equal to wildtype values, and PHM and PAL specific activities in lysates prepared from the sensory-motor cortices of *Pam*^Myh6-cKO/cKO^ mice did not differ from wildtype values.

**Fig.2.**
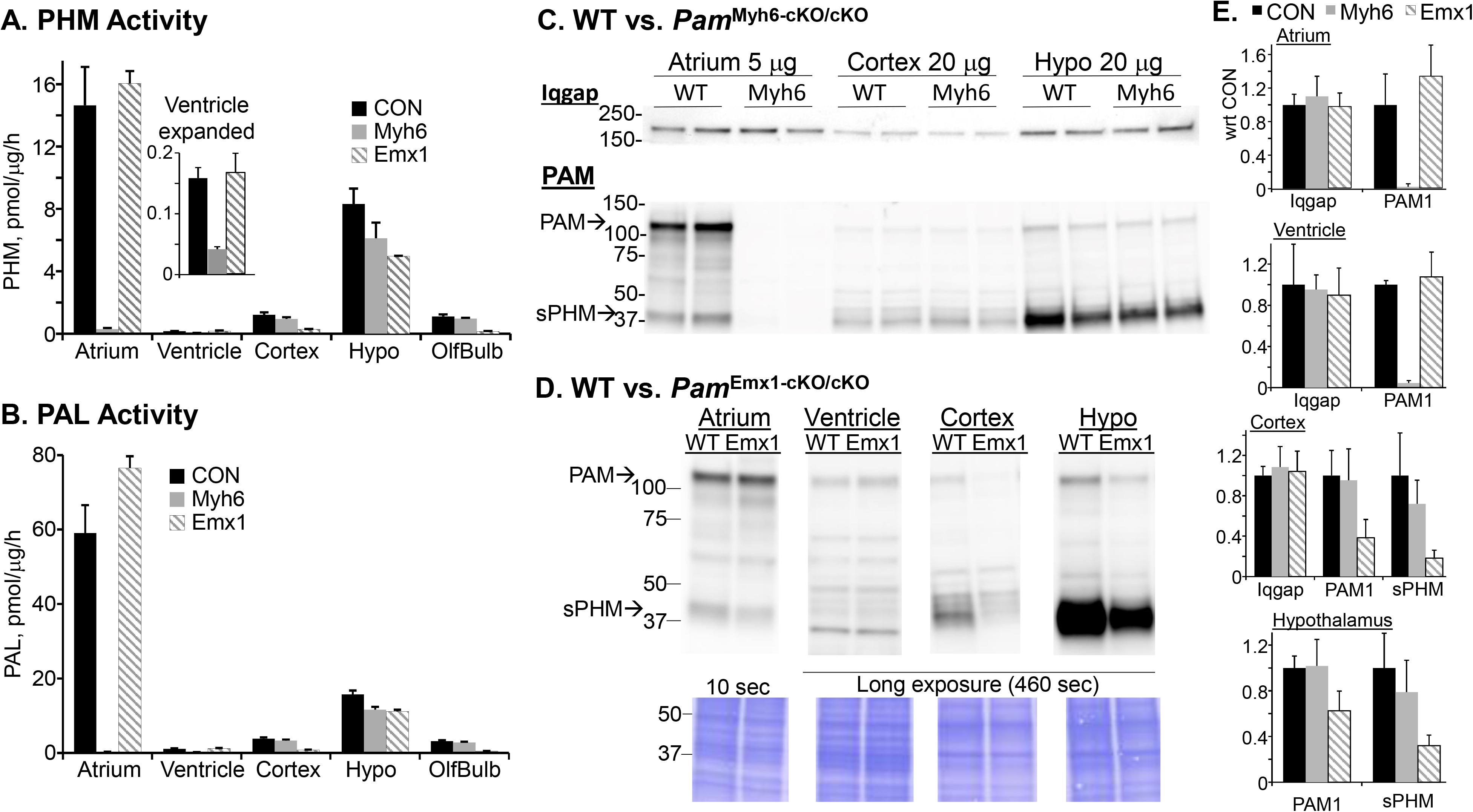
Enzyme assays and Western blots demonstrate tissue-specific elimination of PAM expression. PHM (**A**) and PAL (**B**) activity assays. Data, as pmol product produced/μg lysate protein/hour, are composites of three or more PHM assays and two or more PAL assays of lysates prepared from ≥ 6 mice. **C.** Lysates from CON and *Pam*^Myh6-cKO/cKO^ mice were subjected to Western blot analysis using PAM Exon 16 antibody; the samples shown were analyzed on the same gel, using Iqgap as a loading control. The exposure time shown for the atrial samples was 10 sec, to keep the image in the linear range. **D.** Lysates (20 μg protein, except atrium, 5 μg) prepared from the indicated tissues from WT and *Pam*^Emx1-cKO/cKO^ mice were analyzed in the same manner, with equal loading demonstrated by Coomassie staining. **E.** Western blot data for PAM1 and sPHM were quantified from multiple gel sets; for each gel set, band intensities for knockout tissues were normalized to the corresponding WT band intensity, which was set to 1.00. Data from at least 3 different sample sets were quantified.

Finally, Western blot analyses of lysates prepared from *Pam*^Emx1-cKO/cKO^ and *Pam*^Myh6-cKO/cKO^ mice confirmed the enzyme assay results. In the atrium, intact PAM (110 kDa) was the major product, with smaller amounts of soluble PHM (40 kDa) (**Fig.2C,E**). In both cortex and hypothalamus, soluble PHM was the major product and levels were unaltered in *Pam*^Myh6-cKO/cKO^ mice. In contrast, in the cortex and hypothalamus of *Pam*^Emx1-cKO/cKO^ mice, levels of both intact PAM and soluble PHM were reduced compared to wildtype levels (**Fig.2D,E**). PAM protein levels were unaltered in the atria and ventricles of *Pam*^Emx1-cKO/cKO^ mice.

### Anxiety-like behavior, locomotor response to cocaine and thermoregulation are altered in *Pam*^Emx1-cKO/cKO^ mice

Based on the extensive literature on peptidergic signaling in the nervous system and the phenotypes observed in *Pam*^KO/+^ mice, we first examined *Pam*^Emx1-cKO/cKO^ mice. Previously, *Pam*^KO/+^ mice exhibited a striking increase in anxiety-like behavior compared to wildtype mice (14, 42); this increase was largely ameliorated when the mice were fed a copper supplemented diet. Anxiety-like behavior was tested using the elevated zero maze (**Fig.3A**). Unlike mice with a single allele of *Pam* in all of their tissues (**Fig.3A**, red arrow), *Pam*^Emx1-cKO/cKO^ mice showed an increase in time spent in the open arms of the maze, a response interpreted as a decrease in anxiety-like behavior. Anxiety-like behavior in *Pam*^Emx1-cKO/+^ mice, with a single copy of *Pam* only in Emx1-positive cells, did not differ from control mice (**Fig.3A**).

**Fig.3.**
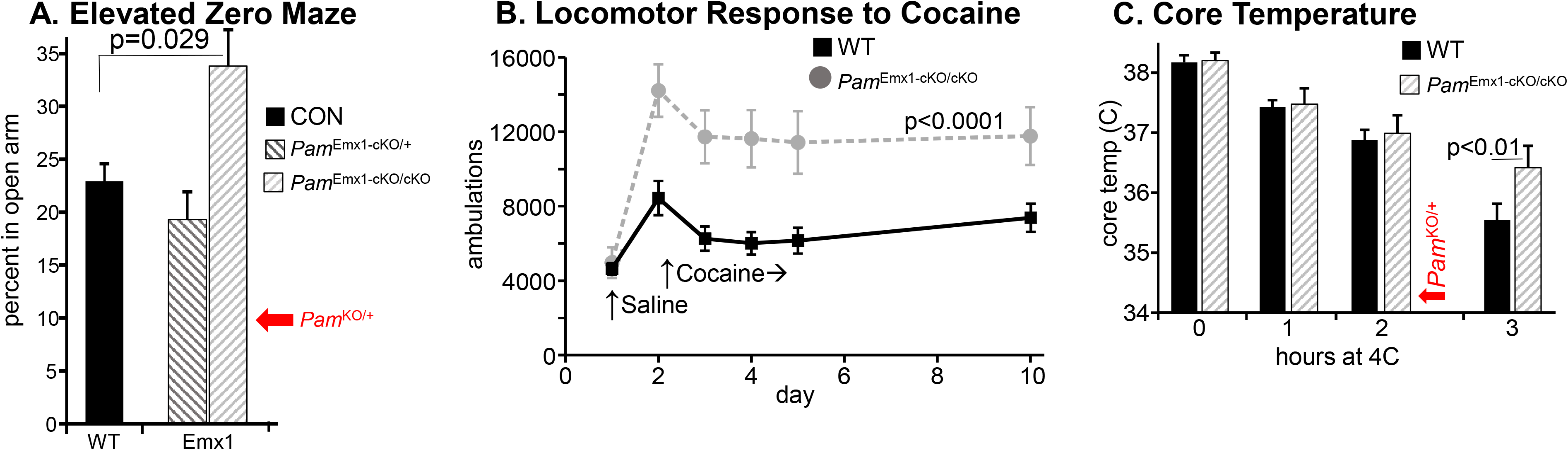
Behavioral/physiological tests of *Pam*^Emx1-cKO/cKO^ mice. **A. Anxiety-like behavior.** Mice ranged from 60 to 88 days old. Control, n = 48; *Pam*^Emx1-cKO/+^, n = 24; *Pam*^Emx1-cKO/cKO^, n = 24. Data for global *Pam* heterozygotes (*Pam*^KO/+^) derived from Emx1-Cre parents (this work) did not differ from data obtained in our previous study (red arrow, *Pam*^KO/+^) (14). **B. Locomotor response to cocaine.** Ambulations were recorded for 45 min after an intraperitoneal injection of saline (Day 1) or cocaine (20 mg/kg on Day 2; 10 mg/kg on Days 3, 4, 5 and 10). Mice ranged from 68 to 151 days old. WT, n = 8; *Pam*^Emx1-cKO/cKO^, n = 8. RM-ANOVA was used to compare knockout to WT. **C. Core body temperature.** Temperature was measured using a rectal thermometer. WT, n = 10; *Pam*^Emx1-cKO/cKO^, n = 20. Mice ranged from 67 to 134 days old. Student’s t-test applied for the 3h data. Previous data from global Pam heterozygotes (*Pam*^KO/+^) are shown by the red arrow (*Pam*^KO/+^) (14).

Studies from many laboratories have revealed a role for amidated peptides in cocaine self-administration and cocaine-seeking behavior during withdrawal (43–48). Neuropeptide Y (NPY), oxytocin and the hypocretins (also called orexins) modulate these behaviors and are inactive if not amidated. We tested the response of *Pam*^Emx1-cKO/cKO^ mice to four injections of cocaine, four days of abstinence and a final injection of cocaine (**Fig.3B**) (34). No difference was observed in the response of wildtype and *Pam*^Emx1-cKO/cKO^ mice to the initial administration of saline (Day 1). *Pam*^Emx1-cKO/cKO^ mice exhibited a much greater response to the initial high dose of cocaine than wildtype mice (Day 2); this increased responsiveness was maintained throughout the entire experimental protocol (RM-ANOVA, p<0.0001).

Thermoregulation was studied because the diminished ability of *Pam*^KO/+^ mice to vasoconstrict peripheral blood vessels led to a marked inability to maintain core body temperature when kept at 4C (14); this deficit forced termination of the previous experiments after 2h in the cold, when body temperature in *Pam*^KO/+^ mice fell to 34C (**Fig.3C**, red arrow). *Pam*^Emx1-cKO/cKO^ mice outperformed control mice when maintained in a 4C environment for a prolonged period of time (**Fig.3C**). In WT mice, body temperature fell significantly between 2 and 3h of cold exposure; *Pam*^Emx1-cKO/cKO^ mice were better able to maintain core body temperature for 3h in a 4C environment (**Fig.3C**). Since *Pam* expression is reduced to half in both excitatory and inhibitory neurons in *Pam*^KO/+^ mice while *Pam*^Emx1-cKO/cKO^ mice lack PAM expression only in excitatory neurons, a difference in their ability to maintain core body temperature was anticipated. Excitatory neurons in the median preoptic area of the hypothalamus promote peripheral dilatation and core heat loss, and produce a number of amidated peptides which strongly affect thermoregulation (49, 50).

### Excitatory neurons in the hippocampus and cortex of *Pam*^Emx1-cKO/cKO^ mice lack PAM

We used a reporter line in which a fluorescent marker, TdTomato, is produced in cells expressing Cre-recombinase to verify the specificity of our knockout strategy at the cellular level (29). To evaluate the overlap between TdTomato-positive cells and PAM-positive cells, PAM was visualized using an antiserum specific for PAM-1 (**Fig.4**); immunocytochemistry controls are presented in **Supplemental Figs.3-6**. As expected (15), TdTomato expression was observed in pyramidal neurons in the CA1 and CA3 regions of the hippocampus in Emx1 Cre-recombinase mice (**Fig.4A,B,C**, yellow arrows). Since the TdTomato reporter is cytosolic, the axons and dendrites of TdTomato-expressing neurons were fluorescent. This was especially apparent in the CA3 region, where a significant fraction of the granule cell mossy fiber endings (mf) innervating the dendrites of the CA3 pyramidal neurons were bright red (**Fig.4C**, mf).

**Fig.4.**
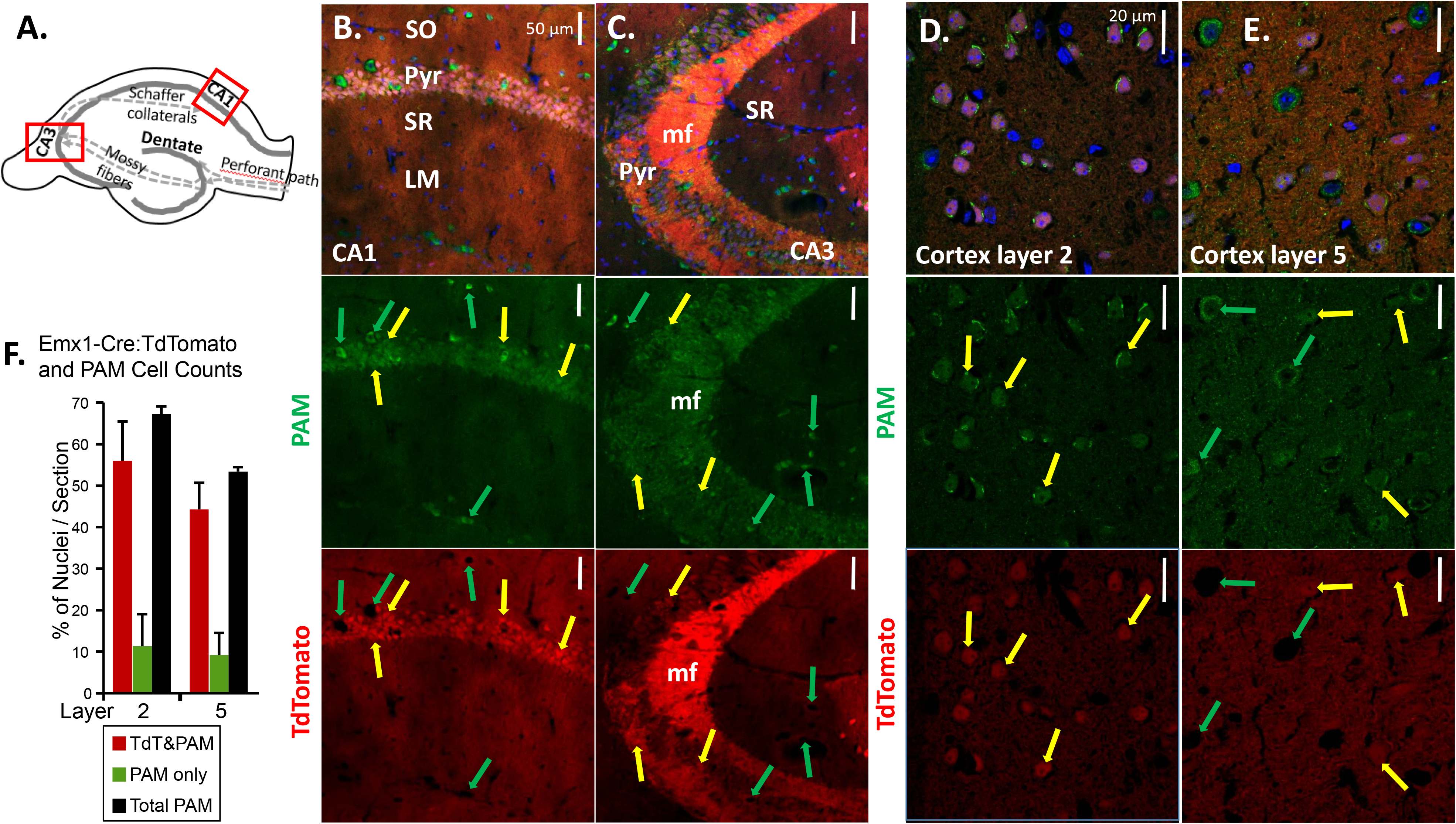
PAM is expressed in all Emx-1 expressing (excitatory) neurons. Adult mice expressing a TdTomato Cre-reporter construct and Emx1-Cre were perfusion fixed; coronal sections were stained for PAM using affinity-purified Exon 16 antibody and AlexaFluor488-tagged secondary antibody (green), while nuclei were visualized using Hoechst stain (blue). Images shown are single plane confocal images. **A.** Diagram of the hippocampus and major pathways; boxed areas indicate the regions of CA1 and CA3 examined. **B.** CA1 region. **C.** CA3 region. **D.** Cortex layer 2. **E.** Cortex layer 5. All TdTomato-positive neurons were also PAM-positive (yellow arrows). A fraction of the PAM-positive neurons did not express TdTomato (green arrows). Abbreviations: stratum oriens, SO; stratum pyramidale, Pyr; stratum radiatum, SR; lacunosum moleculare, LM; mf, mossy fiber. Scale bars, 50 μm (hippocampus) and 20 μm (cortex). **F.** Cell Counts. Cortical images from two mice were quantified (4-5 images per region per Myh6-cKO/cKO animal, 141-213 nuclei per layer per animal): the number of TdTomato/PAM-positive cells and TdTomato-negative/PAM-positive cells is expressed as a percentage of nuclei (Hoechst stained).

Consistent with our previous *in situ* hybridization study (15), the vast majority of pyramidal neurons (Pyr) in both the CA1 and CA3 regions of the hippocampus expressed PAM (AlexaFluor488; green) (**Fig.4B,C**, yellow arrows); PAM staining was also strong in the mossy fibers. Large, strongly PAM-positive neurons that were not TdTomato-positive were scattered throughout the pyramidal cell layer in CA1 and CA3 and in the stratum oriens and stratum lacunosum-moleculare (**Fig.4B,C**, green arrows).

In Layer 2 of the cerebral cortex (**Fig.4D**), all of the TdTomato-expressing neurons (red) were PAM-positive (yellow arrows), suggesting that most of the excitatory neurons in cortical layer 2 could utilize amidated peptides for signaling. PAM-staining in the TdTomato-positive pyramidal projection neurons in cortical layer 5 was less intense, but all of the TdTomato-positive neurons expressed PAM (**Fig.4E**, yellow arrows). The layer 5 neurons with the most intense PAM signal were not TdTomato-positive, suggesting that they were interneurons (**Fig.4E**, green arrows).

The number of neurons in layers 2 and 5 expressing TdTomato and stained for PAM and the number of PAM-positive neurons not expressing TdTomato was compared to the total number of nuclei (**Fig.4F**). In both layers, the number of neurons expressing PAM, but not expressing TdTomato was approximately one fifth of the number of neurons expressing both PAM and TdTomato.

### PAM is highly expressed in GAD67-positive interneurons

Given the importance of excitatory/inhibitory balance in nervous system function, we wanted to identify PAM-positive inhibitory neurons; to do so, coronal sections from control and *Pam*^Emx1-cKO/cKO^ mice were stained simultaneously for PAM (green) and glutamic acid decarboxylase 67 (GAD67) (red) (**Fig.5**). The strongly PAM-positive neurons seen in the stratum oriens and stratum lacunosum moleculare of the CA3 region in both control and *Pam*^Emx1-cKO/cKO^ mice were largely GAD-positive (Fig.5A, yellow arrows).

**Fig.5.**
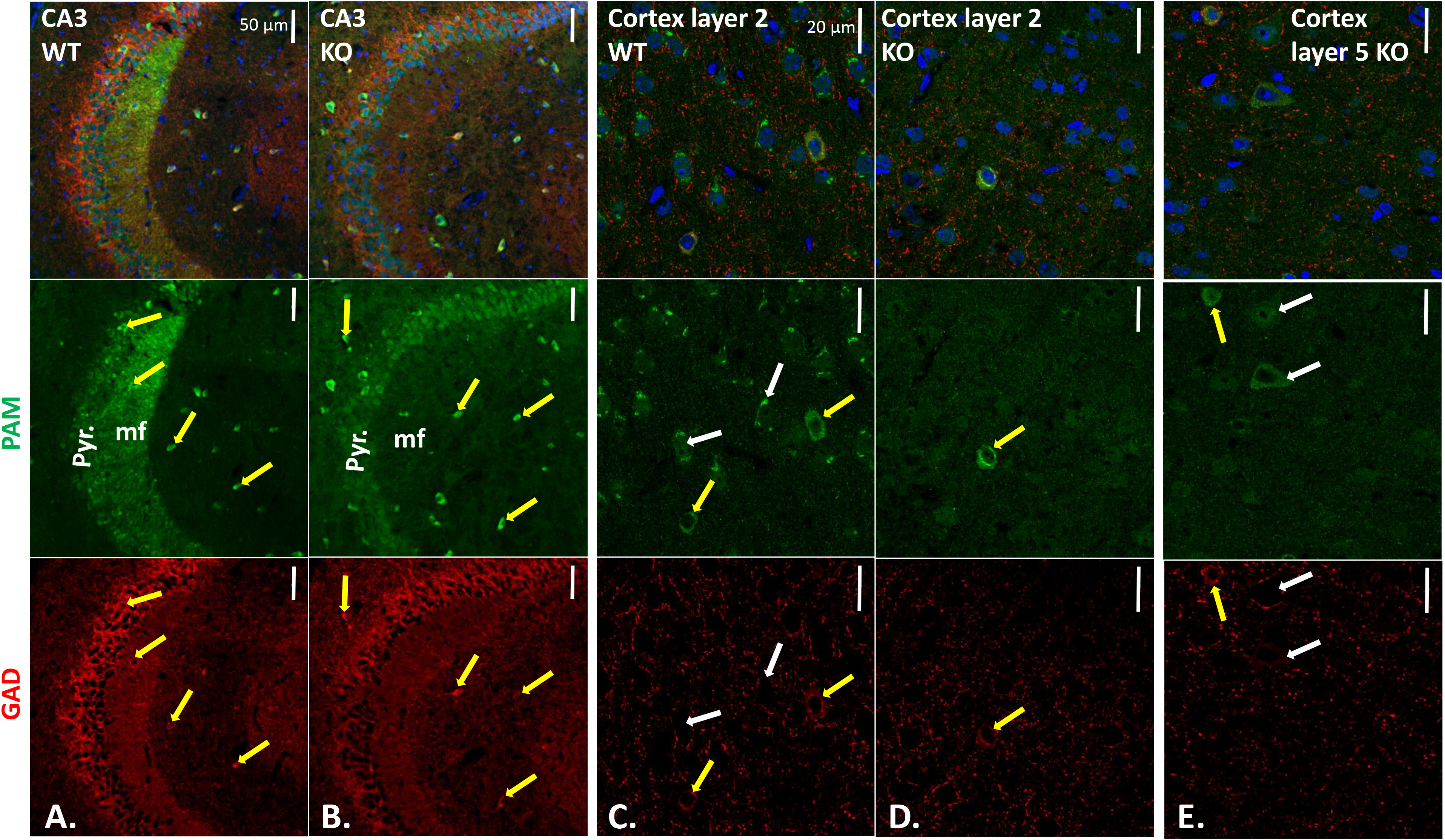
Many of the PAM-positive neurons that remain in the hippocampus and cortex of *Pam*^Emx1-cKO/cKO^ mice express GAD. Sections stained with affinity-purified PAM-Exon 16 antibody plus AlexaFluor488-antibody to rabbit IgG (green), mouse monoclonal antibody to GAD67 plus Cy3-antibody to mouse IgG (red), and Hoechst nuclear stain (blue). CA3 regions of WT (**A.**) and KO (**B.**) hippocampus; yellow arrows mark neurons stained for PAM and GAD67. WT (**C.**) and KO (**D.**) layer 2 and KO layer 5 (**E.**) visualized as described for hippocampus. Scale bars, 50 μm (hippocampus) and 20 μm (cortex). In KO layer 2, all GAD-positive neurons were PAM-positive; in KO layer 5, some PAM-positive neurons were not GAD-positive (white arrows). Quantification is in **Supplemental Table 2**.

Strikingly, the PAM-positive mossy fibers observed in the wildtype CA3 region were not seen in the *Pam*^Emx1-cKO/cKO^ brain (**Fig.5B**). In wildtype mice, neuropeptide Y (NPY), an amidated peptide, is highly expressed in mossy fibers (30). The diffuse nature of the AlexaFluor488 signal observed in pyramidal neurons in the *Pam*^Emx1-cKO/cKO^ mice (**Fig.5B**) resembled the background observed when PAM antibody was replaced by non-immune rabbit immunoglobulin or blocked with antigenic peptide (**Suppl.Figs.4-7**).

In cortical layer 2 (**Fig.5C, D**), very few PAM-positive neurons were identified in *Pam*^Emx1-cKO/cKO^ mice; almost all of the remaining PAM-positive neurons expressed GAD67 (inhibitory neurons) (**Fig.5D**, yellow arrow; **Suppl.Fig.6,7**). The PAM-positive neurons in cortical layer 5 of *Pam*^Emx1-cKO/cKO^ mice (**Fig.5E**, yellow arrow), were almost always GAD67-positive (inhibitory) (**Suppl.Fig.6,7**). In the cerebral cortices of WT mice, there were PAM-positive neurons not positive for GAD, as expected from the majority of PAM-positive neurons that were Emx1-positive (**Fig.4**) (white arrows). In addition, in *Pam*^Emx1-cKO/cKO^ mice, there were PAM-positive neurons that were not GAD-positive (**Fig.5E**, white arrows). The decreases in PAM-positive neuron numbers in cortical layers 2 and 5 in *Pam*^Emx1-cKO/cKO^ mice (L2: 56%; L5: 39%) closely matched the number of neurons expressing both PAM and TdTomato in the Emx1-Cre reporter mice (L2: 56.0 ± 9.5%; L5: 44.3 ± 6.4%) (summarized in **Supplemental Table 2**).

### Anxiety-like behavior and thermoregulation are altered in *Pam*^Myh6-cKO/cKO^ mice

Although atrial levels of PAM are at least 10-fold higher than cortical levels (18) (**Fig.2**), neither proANP nor pro-brain natriuretic peptide (pro-BNP) is amidated. The tests used to characterize *Pam*^Emx1-cKO/cKO^ mice (**Fig.3**) were also used to evaluate *Pam*^Myh6-cKO/cKO^ mice (**Fig.6**). When tested for anxiety-like behavior, the *Pam*^Myh6-cKO/cKO^ mice responded in much the same way as *Pam*^KO/+^ mice; time spent in the open arms decreased to about half of the value observed in control mice (**Fig.6A**). The anxiety-like behavior of *Pam*^Myh6-cKO/+^ mice did not differ from that of control mice. When tested using the cocaine injection paradigm described in **Fig.3B**, *Pam*^Myh6-cKO/cKO^ mice showed an enhanced response only on Day 2 (p<0.02); unlike *Pam*^Emx1-cKO/cKO^ mice, the elevated response was not sustained (**Fig.6B**). Like *Pam*^Emx1-cKO/cKO^ mice, *Pam*^Myh6-cKO/cKO^ mice were better able than wildtype mice to maintain core body temperature for 3h in a 4C environment (**Fig.6C**).

**Fig.6.**
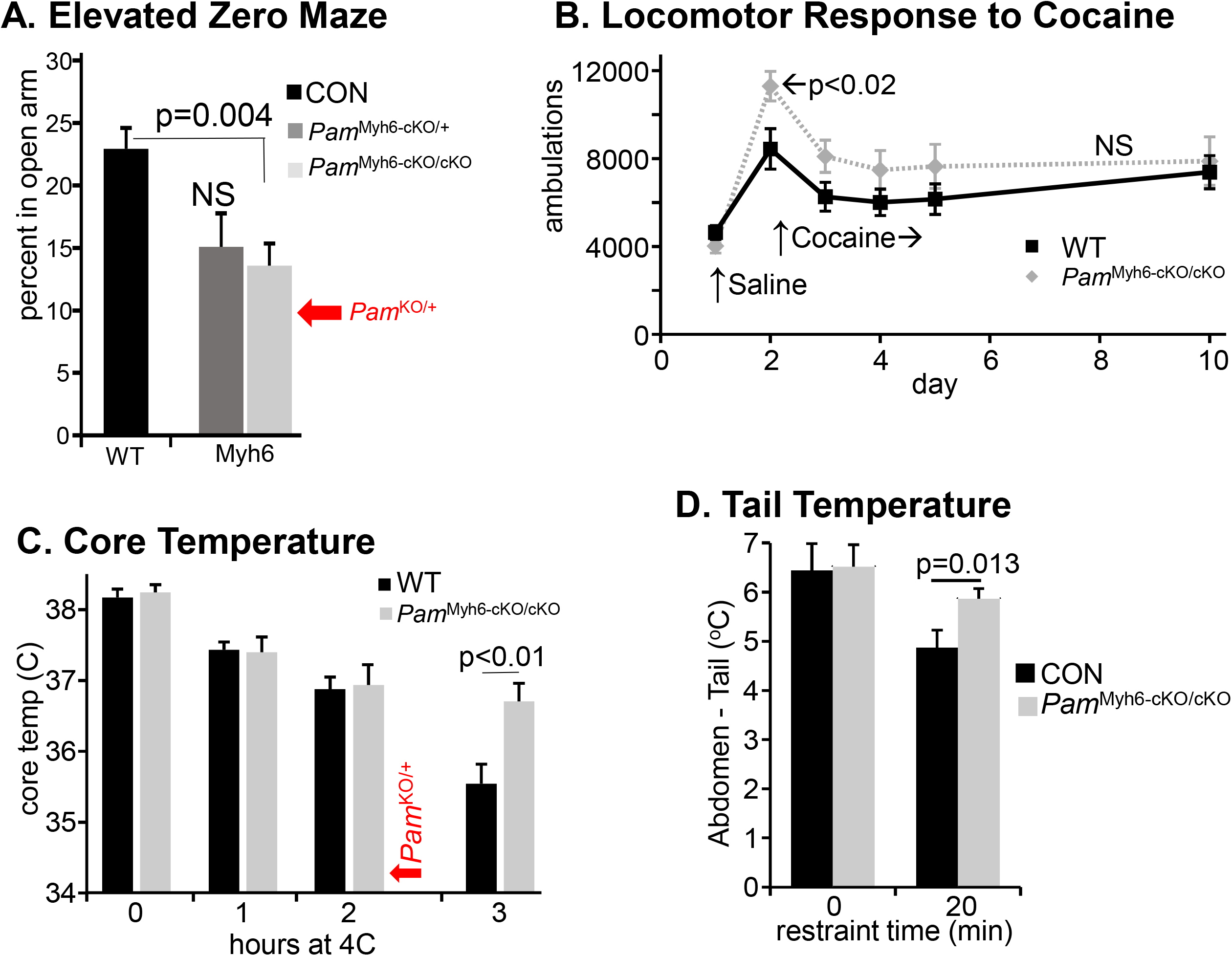
Behavioral/physiological tests of *Pam*^Myh6-cKO/cKO^ mice. *Pam*^Myh6-cKO/cKO^ mice were tested along with the mice reported in **Fig.3**; control data are replicated here. **A. Anxiety-like behavior.** Mice ranged from to 88 days old. Control, n = 48; *Pam*^Myh6-cKO/+^, n = 20; *Pam*^Myh6-cKO/cKO^, n = 30. Data obtained previously for *Pam*^KO/+^ are shown (red arrow) (14). **B. Locomotor response to cocaine.** Ambulations after an intraperitoneal injection of saline or cocaine were recorded as described in **Fig.3**. Mice ranged from 61 to 114 days old. WT, n = 8; *Pam*^Myh6-cKO/cKO^, n = 12. RM-ANOVA was used to compare knockout to Control. **C. Core body temperature.** WT, n = 10; *Pam*^Myh6-cKO/cKO^, n = 10. Mice ranged from 67 to 134 days old. Student’s t-test applied for the 3h data. Previous data for *Pam*^KO/+^ are shown by the red arrow (14). **D. Abdominal and tail temperatures during 20 min restraint in an unfamiliar environment.** Abdominal temperatures were determined at the beginning and end of 20 min restraint, and tail temperatures were determined throughout the experiment. The difference (Abdomen minus tail) at 0 and 20 min is shown. CON, n=20; *Pam*^Myh6-cKO/cKO^, n = 23.

The marked improvement in the ability of the *Pam*^Myh6-cKO/cKO^ mice to maintain core body temperature for 3h suggested that these mice had an enhanced coping mechanism whereby they could redirect peripheral blood flow to the core (35, 36). A common test for this ability is use of a heat-sensitive probe to monitor tail temperature while the animal is kept in an unaccustomed restraint device for 20 min (35, 36); abdominal temperature is measured with the same probe before and after the 20 min restraint, and the temperature differential (abdomen minus tail) is reported (**Fig.6D**). Measured in this manner, abdominal temperature was the same in *Pam*^Myh6-cKO/cKO^ and WT mice both before and after restraint. The tail temperature differential was significantly greater in *Pam*^Myh6-cKO/cKO^ mice than in WT mice after 20 min of restraint, indicating an enhanced ability of *Pam*^Myh6-cKO/cKO^ mice to constrict tail blood flow.

### Atrial levels of pro-atrial natriuretic peptide (proANP) decline in *Pam*^Myh6-cKO/cKO^ mice

As expected from the literature (19, 20) and consistent with the dramatically reduced levels of PAM protein and enzyme activity measured in the atria and ventricles of *Pam*^Myh6-cKO/cKO^ mice (**Fig.2**), TdTomato was uniformly expressed in atrial and ventricular cardiomyocytes when driven by the Myh6 promoter (**Suppl.Fig.7**).

PAM is the major protein in the membranes of atrial secretory granules, which store proANP and proBNP (7, 51–53). PAM interacts directly with proANP, which is not cleaved until the time of secretion Myh6-cKO/cKO Myh6-cKO/cKO Myh6-cKO/cKO (25, 54). Lysates prepared from WT and *Pam*^Myh6-cKO/cKO^ atria using 1% deoxycholate to ensure better protein solubilization were subjected to western blot analysis (**Fig.7A**). As assessed using antibodies to PAM Exon 16 (JH629) and to the cytoplasmic domain of PAM (C-Stop), PAM levels in *Pam*^Myh6-cKO/cKO^ atria were negligible. ANP levels were assessed using an antibody specific for the N-terminal region of proANP and an antibody specific for a 15-residue peptide contained within mature ANP, a 28-residue peptide. Both antibodies revealed a dramatic drop in proANP levels in *Pam*^Myh6-cKO/cKO^ atria (**Fig.7A**).

**Fig.7.**
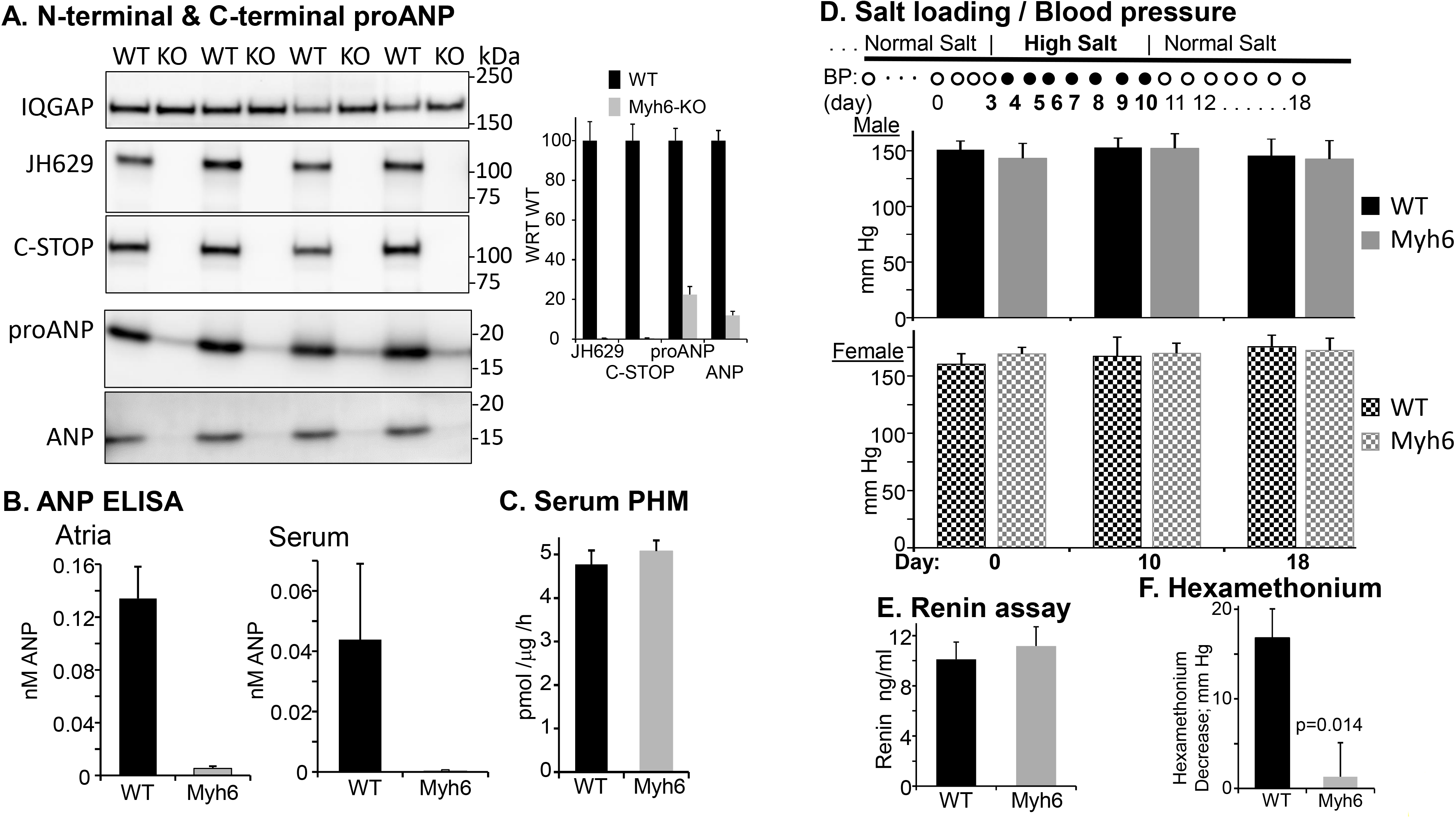
Alterations in proANP and blood pressure regulation in *Pam*^Myh6-cKO/cKO^ mice. **A.** RIPA lysates of four pairs of WT and *Pam*^Myh6-cKO/cKO^ atria (1 μg protein/lane) were fractionated by SDS-PAGE and analyzed for PAM (PAM-1 specific antibody JH629; PAM cytosolic domain antibody, C-Stop) and proANP; the gel that was analyzed using the ANP-specific antibody had 10 μg protein/lane. Graph shows quantification, relative to WT. **B.** ANP ELISAs confirmed the drop in ANP levels in atrial extracts and trunk blood (N=10 for serum, 14 for tissue). **C.** PHM assays on serum samples from WT and *Pam*^Myh6-cKO/cKO^ mice (WT n=3; *Pam*^Myh6-cKO/cKO^, n=3). Similar results were obtained in a second assay comparing these mice, as well as sera from WT and *Pam*^Emx1-cKO/cKO^ mice (not shown). **D. High Salt Diet.** Tail cuff blood pressure was recorded for WT mice (5 male, 3 female) and *Pam*^Myh6-cKO/cKO^ mice (3 male, 5 female). Mice were maintained for >2 months on a normal salt diet (0.49% NaCl), 7 days on a solid diet including 8% NaCl (High Salt), during which body weight and food consumption remained constant, followed by an additional week on the normal salt diet. Mice were 79-101 days old. The experiment was repeated with WT mice (2 male, 2 female) and *Pam*^Myh6-cKO/cKO^ mice (2 male, 2 female) with the same results. **E.** Serum renin was measured by ELISA, with 16 WT and 16 *Pam*^Myh6-cKO/cKO^ samples. **F.** Tail cuff blood pressure was recorded 10 min after an injection of isotonic saline (Day 1) and after an injection of hexamethonium (Day 2) (38, 39). The average difference between the Day 1 and Day 2 data for each mouse is plotted. Fourteen WT mice and 15 *Pam*^Myh6-cKO/cKO^ mice, 59-102 days old.

To determine whether the decline in proANP levels in the atria of *Pam*^Myh6-cKO/cKO^ mice was accompanied by altered circulating levels of ANP, we utilized an ELISA to quantify atrial and serum levels of ANP in *Pam*^Myh6-cKO/cKO^ and WT mice (**Fig.7B**). Levels of ANP in the atria and sera of *Pam*^Myh6-cKO/cKO^ mice were dramatically lower than in control mice. Although PAM expression in the atrium exceeds levels in any other tissue examined, levels of serum PHM activity were unaltered in *Pam*^Myh6-cKO/cKO^ mice (Fig.7C), eliminating the atrium as a major source of serum PAM.

### Blood pressure regulation is altered in *Pam*^Myh6-cKO/cKO^ mice

ANP has been referred to as a natural antihypertensive agent (55); along with several other peptides, ANP participates in the control of blood pressure and the ability of mammals to respond to changes in dietary salinity and cardiovascular fluid volume (38, 56–58). Since circulating levels of ANP were undetectable in *Pam*^Myh6-cKO/cKO^ mice, their ability to regulate blood pressure in response to a change in dietary salt load was examined. When fed a normal salt diet (0.49% NaCl) for two months, no differences in blood pressure were observed between control and *Pam*^Myh6-cKO/cKO^ mice (**Fig.7D**, **Day 0**); blood pressure was consistently slightly higher in female mice than in male mice, irrespective of PAM genotype.

This result suggested that other systems might be compensating for the low levels of ANP in *Pam*^Myh6-cKO/cKO^ mice. The renin-angiotensin system increases blood pressure by stimulating the conversion of angiotensinogen to angiotensin I; renin secretion is stimulated by low blood sodium and sympathetic input, and suppressed by ANP (59). However, assays for serum renin levels in WT and *Pam*^Myh6-cKO/cKO^ mice on a normal salt diet revealed no difference (**Fig.7E**).

The WT and *Pam*^Myh6-cKO/cKO^ mice, whose blood pressure was monitored after two months on a normal salt diet, were fed a high salt diet (8% NaCl) for a week (**Fig.7D;** days 4-10). No changes in blood pressure or body weight were observed, despite the low levels of ANP found in *Pam*^Myh6-cKO/cKO^ mice. These mice were then returned to the normal salt diet (days 11-18), again with no changes observed in blood pressure.

The sympathetic system, a major ANP target (38, 60), plays an essential role in blood pressure control. Hexamethonium, a ganglionic blocker that eliminates all presynaptic sympathetic ganglion input, was used to test the hypothesis that sympathetic tone differed in WT and *Pam*^Myh6-cKO/cKO^ mice (38, 39, 61). As expected, control mice exhibited an approximately 20 mm Hg decrease in arterial blood pressure 10 min after an injection of hexamethonium (**Fig.7F**). In contrast, blood pressure in *Pam*^Myh6-cKO/cKO^ mice was unaltered after hexamethonium injection, revealing a striking absence of sympathetic tone.

### The transcriptome is altered in *Pam*^Myh6-cKO/cKO^ atrium

The homogeneity of atrial tissue, lack of a major amidated product despite high levels of *Pam* expression and changes observed in the *Pam*^Myh6-cKO/cKO^ mice led us to ask whether changes in gene expression might contribute to the phenotypes observed. Alterations in the atrial transcriptome could reflect the inability of these cells to produce sfCD (soluble fragment of the PAM cytosolic domain, which translocates to the nucleus) (26), a direct response to the diminished levels of ANP or compensatory responses by the many systems involved in regulating these fundamental processes. Atrial RNA was prepared from WT and *Pam*^Myh6-cKO/cKO^ mice and sequenced (**Fig.8A**; **Suppl.Table3**).

**Fig.8.**
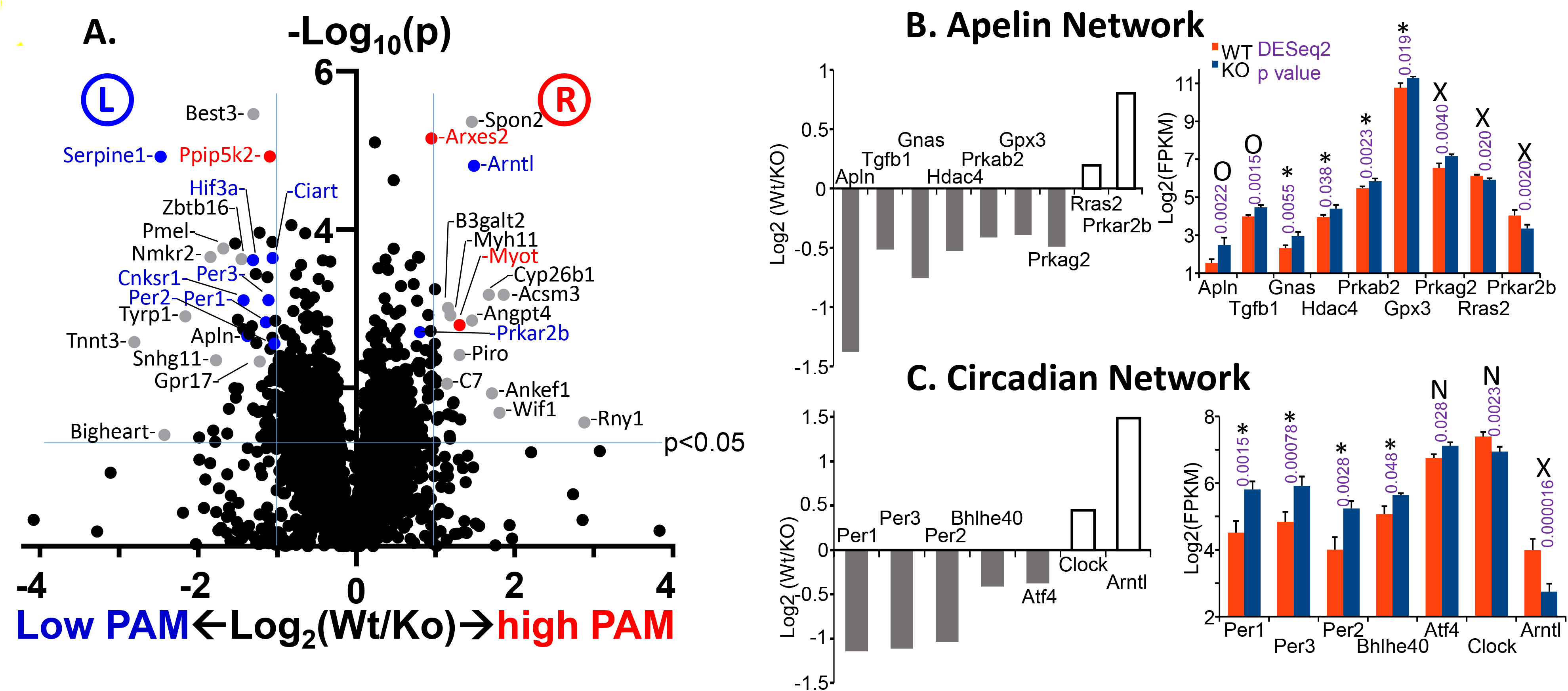
RNAseq of Atrial RNA. **A.** Volcano plot of all transcripts with FPKM>1, comparing WT and *Pam*^Myh6-cKO/cKO^ samples; the horizontal blue line indicates the cutoff for DESeq2 significance (p<0.05), while the vertical blue lines indicate transcripts changed at least 2-fold up or down, comparing WT and *Pam*^Myh6-cKO/cKO^ samples. X-axis is Log2 of the ratio (WT/KO); Y-axis is −Log10 of the DESeq2 p value (up is more significant). Transcripts with blue dots were decreased when PAM was induced in pituitary cells in our previous study (10), while red dots indicate transcripts increased when PAM was induced. Grey dots indicate transcripts expressed at low levels in pituitary cells (FPKM<3). **B,C.** From Ingenuity Pathway Analysis, two of the most significant networks were centered on the Apelin peptide and on circadian rhythms. The y-axes of the bar graphs on the left show Log_2_ of the Wt/KO ratio. The y-axes of the bar graphs on the right show Log_2_ of the FPKM values for each transcript in Wt and KO, along with the DESeq2 p value. Code: *, significant in pituitary cells (10); N, no change in pituitary cells; 0, not expressed above FPKM=2 in pituitary cells; X, opposite response in pituitary cells.

DESeq2 analysis of differential expression was performed on the counts data, yielding 1063 transcripts with p<0.05; when outliers were eliminated (34), the number of differentially expressed transcripts dropped to 555. When depicted as a volcano plot, the vast majority of over 14,000 transcripts with average FPKM>1 were clustered near the origin, with a WT/KO ratio within the 0.5-2.0 range (Log2[ratio] −1 to +1) and a DESeq2 p value > 0.05.

The 555 significantly altered transcripts were subjected to Ingenuity Pathway Analysis (IPA) (**Fig.8B,C**; **Suppl.Fig.8**) (34). One of the most significantly regulated networks identified by IPA was the Apelin network. Apelin (Gene ID 8862) is an adipokine made in the atrium whose FPKM level rose 2.6-fold in *Pam*^Myh6-cKO/cKO^ mice compared to WT. Apelin is used clinically to treat insulin resistance and hypertension (62). Another major network includes a set of transcripts involved in circadian rhythms (three Period genes, *Clock*, *Bhlhe40* and *Arntl*). Bhlhe40 is a basic-helix-loop-helix protein which binds to the promoter of *Per1* to repress Clock/Arntl activation of *Per1*; Bhlhe40 levels vary directly with blood pressure in Clock knockout mice (63). Three more significantly regulated networks identified by IPA are depicted in **Suppl.Fig.8**. The tight interconnections of these five predicted networks are striking, with several transcripts appearing in two or three of the five networks: Apln, Apold1, Gadd45b, Bhlhe40, and Per1.

Many of the differentially expressed atrial transcripts were also identified as *Pam*-regulated genes in pituitary corticotrope cells (10). Strikingly, 20 of the 38 atrial transcripts in the five most significantly regulated networks responded to changes in Pam expression in the same manner in these two very different systems (marked with asterisks [*]). For example, expression of *Per1* increased more than 2-fold in *Pam*^Myh6-cKO/cKO^ mice compared to WT and decreased to 60% when *Pam* expression was increased in pituitary cells (10), suggesting that the change may be a direct response to altered PAM levels. Expression of 10 atrial transcripts (marked with ‘X’) decreased with loss of atrial PAM and also decreased in pituitary cells when PAM was increased (10), suggesting that these transcripts are not responding directly to the PAM protein but rather to some other compensatory changes or differences between the two tissues. Five of the 38 atrial transcript are not expressed at significant levels in pituitary cells (marked with ‘0’).

## DISCUSSION

### Rationale

Peptidergic signaling in species as diverse as *Trichoplax*, *Drosophila* and human shares many common features (8). The preproproteins synthesized into the lumen of the ER undergo a series of post-translational modifications that generate a wide variety of products. In many cases these modifications are tissue-specific and generate multiple products that can interact with different receptors. The receptors, which are almost always G protein coupled receptors, may be located nearby or far away. Peptides play essential roles in the organism’s response to nutrients and to noxious stimuli, function to distinguish self from non-self, participate in sexual reproduction, and control motility through their effects on ciliary beating and muscle contractility. Peptide expression is regulated at multiple levels, with changes in transcription, translation and post-translational processing as well as acute effects on peptide release. This complexity has made it difficult to utilize specific peptides in the development of therapeutics.

To test the utility of *Pam*^cKO/cKO^ mice in revealing functionally significant effects of peptidergic signaling, we selected two Cre drivers. Based on the widespread expression of PAM throughout the nervous system (**Figs.4**,**5**; (15)), we utilized the Emx1-Cre mouse. A proper balance of excitatory and inhibitory inputs is critical to the functioning of many circuits and we hypothesized that loss of PAM in excitatory neurons would disrupt this balance. Eliminating the expression of vasoactive intestinal polypeptide, an amidated peptide, in inhibitory interneurons in the auditory cortex, alters acoustic gain (64). In the feeding-satiety network, different sets of neurons, many producing amidated peptides, battle for dominance (65). The Myh6-Cre mouse was selected to gain insight into the roles played by PAM in atrial myocytes, which produce massive amounts of PAM (18, 51, 52), but are not known to produce amidated peptides (**Fig.2**). Atrial myocytes store intact proANP in secretory granules whose major membrane protein is PAM (51, 52).

Immunostaining readily identified PAM in every TdTomato-positive neuron (**Fig.4**). This observation is consistent with the four-fold drop in PHM and PAL activity observed in the cortices of the *Pam*^Emx1-cKO/cKO^ mice. A subset of the GABA-positive neurons expressed PAM; in general, PAM staining in GABAergic neurons was more intense than in other neurons. Mice entirely lacking the ability to express PAM did not survive beyond E14.5 (11), with massive edema accompanied by a poorly formed vasculature. Surprisingly, mice lacking the ability to express PAM only in their cardiomyocytes did not exhibit these defects. *Pam*^Myh6-cKO/cKO^ mice grew normally, performed as well as WT mice on the RotaRod and in the Open Field, and outperformed wildtype mice in thermoregulation tests.

### Anxiety-like behavior

The amygdala plays a crucial role in anxiety (66). Anxiety-like behavior increased in *Pam*^KO/+^ mice, with reduced levels of PAM expression in both excitatory and inhibitory neurons (14). In contrast, eliminating *Pam* expression only in excitatory neurons resulted in a decrease in anxiety-like behavior (**Fig.3A**). Treatment of *Pam*^KO/+^ mice with diazepam, a GABA-A receptor agonist, normalized their anxiety-like behavior, suggesting that it resulted from limited inhibitory tone (14, 42). Ifenprodil, which blocks a subtype of excitatory glutamate receptors, has anxiolytic actions in wildtype mice (67). If the net effect of the amidated peptides expressed in excitatory neurons is to facilitate the actions of glutamate, eliminating their expression in *Pam*^Emx1-cKO/cKO^ mice would be expected to reduce anxiety-like behavior (**Fig.3A**). The deficits observed in the *Pam*^Emx1-cKO/cKO^ mice indicate that further study is warranted, despite the large number of neuropeptides and receptors involved, the complexity of the circuits and the lack of specific pharmacological blockers.

The increase in anxiety-like behavior observed in *Pam*^*KO/+*^ mice was mimicked in the atrium-specific total knockout *Pam*^Myh6-cKO/cKO^ mice (**Fig.6A**). The reduction in circulating ANP in *Pam*^Myh6-cKO/cKO^ mice may contribute to this phenotype. There is a well-recognized correlation between high anxiety and low circulating ANP levels in human cardiovascular patients (68); a subset of patients with low anxiety have high circulating levels of the N-terminal pro-ANP (69). It is striking that a number of transcripts which could adversely affect cardiac function were altered in *Pam*^Myh6-cKO/cKO^ mice, potentially contributing to anxiety: increased Hdac4, promoting myocardial ischemia; increased Prkag2, lowering heart rate; increased Usp2, blunting mineralocorticoid responsiveness and decreasing retention of high salt; decreased Ano1 (a Ca^++^ activated Cl^−^ channel) and increased Cacna1a (Ca^2+^ channel), both potentially increasing the strength of heart contraction.

### Response to cocaine

The locomotor response to cocaine involves altered signaling in the nucleus accumbens, with established roles for dopamine, glutamate, and amidated neuropeptides such as NPY, oxytocin and the hypocretins (Hcrt) (70–72). Activation of the NPY Y1 receptor in the amygdala increases self-administration of psychostimulants such as cocaine, while activation of the NPY Y2 receptor inhibits self-administration (43, 48). Hcrt1 and Hcrt2 are amidated peptides derived from adjacent regions of the Hcrt precursor. The Hcrtr1 receptor binds Hcrt1 much better than Hcrt2 (44) and selective inhibitors of Hcrtr1 receptors are in clinical trials to treat cocaine addiction (44). Systemic delivery of oxytocin reduces cocaine self-administration (47) (also in clinical trials). The anti-addictive actions of oxytocin, NPY (at the Y2 receptor) and hypocretins (at Hcrtr2) could be missing in *Pam*^Emx1-cKO/cKO^ mice (**Fig.3B**) but not in *Pam*^Myh6-cKO/cKO^ mice (**Fig.6B**).

### Thermoregulation

*Pam*^Emx1-cKO/cKO^ mice withstand cold exposure better than wildtype mice (**Fig.3C**). The preoptic area of the hypothalamus is the central control region for core body temperature (50). Glutamatergic (excitatory, Emx1-expressing) neurons drive peripheral dilatation and severe hypothermia (49), while GABAergic neurons have less effect (73). Excitatory neurons in the preoptic area express many amidated neuropeptides: PACAP, CCK, CRH, oxytocin, tachykinin1 and GnRH (50, 74). Importantly, ablation of medial preoptic excitatory neurons causes dramatic hyperthermia (49, 75); deletion of PAM in these neurons (as for ablation) could produce peripheral vasoconstriction and core heat gain.

*Pam*^Myh6-cKO/cKO^ mice also withstand cold exposure better than wildtype mice (**Fig.6C**). The RNAseq results provide potential explanations for this, with markedly increased transcript levels of Gadd45g (Growth arrest and DNA-damage-inducible 45 gamma; also called Ddit2) and decreased Prkar2b (Protein kinase, cAMP dependent regulatory, type 2 beta). Both changes could increase the expression of uncoupling protein 1 (Ucp1), which is thermogenic, increasing metabolic rate. In addition, *Pam*^Myh6-cKO/cKO^ mice were more capable than wildtype mice of decreasing heat loss from the periphery (tail, limbs) under stress (restraint or coldroom) (**Fig.6D**).

### Blood pressure

*Pam*^Myh6-cKO/cKO^ mice showed no increase in blood pressure in response to salt loading (8% NaCl diet), matching wildtype mice (**Fig.7D**). Use of hexamethonium revealed that 20 mm Hg of the wildtype mouse blood pressure was due to resting sympathetic nervous tone, while *Pam*^Myh6-cKO/cKO^ mice lacked any resting sympathetic tone (**Fig.7F**). *Pam*^Myh6-cKO/cKO^ mice have negligible circulating ANP (**Fig.7B**), as in ANP and Corin knockout mice, which exhibit salt-induced hypertension of 15-30 mm Hg (28, 38, 76). ANP and BNP are released from the atrium in response to cardiac distension (38, 56). Wildtype mice fed solid diets containing 0.49% (normal) and 8% (high) NaCl show no detectable elevation in arterial blood pressure in most studies (28, 38, 76); only 25% of normotensive humans are salt sensitive, by the American Heart Association (77).

ANP, vasopressin, angiotensin II, bradykinin and neuropeptide Y are the dominant peptides controlling blood pressure responses to dietary salt and altered cardiovascular volume (38, 56, 58). Only vasopressin and NPY require amidation for bioactivity, and both elevate blood pressure, so loss of amidation could decrease blood pressure. ANP works in opposition to the sympathetic system, attenuating vascular tone, inhibiting vasopressin release, reducing renin release and decreasing angiotensinogen conversion into angiotensin (38, 56, 58). Since circulating renin levels were unaltered in *Pam*^Myh6-cKO/cKO^ mice (**Fig.7E**), we hypothesize that lack of sympathetic tone (**Fig.7F**) counterbalances loss of circulating ANP (**Fig.7C**).

### Transcript changes in atria of *Pam*^Myh6-cKO/cKO^ mice

Levels of about 550 atrial transcripts were significantly altered in *Pam*^Myh6-cKO/cKO^ mice (**Fig.8**, **Suppl.Fig.8, Suppl.Tables 3,4**), just over 2% of the genome. Altered transcript levels may underlie some of the physiological changes seen in *Pam*^Myh6-cKO/cKO^ mice, such as increased anxiety, heightened ability to withstand a cold challenge, and lack of increased blood pressure on an 8% NaCl diet (**Figs.6**,**7**). One interesting transcript increased in *Pam*^Myh6-cKO/cKO^ mice encodes apelin, a secreted protein. Signal peptide removal yields proapelin, a 55 amino acid protein; the 13 amino acid COOH-terminal peptide that binds to the APJ receptor is preceded by a furin-like cleavage site expected to generate an amidated 34 amino acid product. After experimentally-induced pressure overload due to aortic banding, *Apln* knockout mice exhibit severely reduced cardiac contractility (78).

The generation of a PAM-derived COOH-terminal fragment called sfCD, which enters the nucleus (26), led to studies on the effects of PAM on gene expression. We know from our studies in a doxycycline-inducible corticotrope tumor cell line that a 2-day 100-fold increase in PAM expression (up close to the level in the adult atrium) altered the expression of a few hundred genes (10). The altered atrial gene expression observed here could reflect the absence of PAM itself or a compensatory response to the consequences of lacking PAM expression since conception. It is particularly interesting to note the large number of transcripts which change in response to altered PAM expression in both pituitary cells and atrium. Of the 100 transcripts most altered in the atrium by PAM deletion since conception (quadrants L and R in **Fig.8A**, with DESeq2 p<0.05 and more than a 2-fold change), over a third were also responsive to a 48 h increase in PAM expression in pituitary cells (10). A majority of these changes correlated with PAM levels in these two very different model systems.

## Acknowledgments

We thank Dr. Nils Bäck for the broad perspective he provided as he offered comments on this manuscript and members of the Neuropeptide Lab for many helpful discussions. This work was supported by NIH Grants R01DK032948 and R01DK032949, the Janice and Rodney Reynolds Endowment and the Daniel Schwartzberg Fund.

