## Supplemental Text for "Identifying Novel Roles for Peptidergic Signaling in Mice"

**Supplemental Methods**

**Genotyping the mice**

Polymerase chain reaction (PCR) screening of DNA prepared from ear punches or tail clips was performed using reactions with three primers (**Suppl.Fig.1**). Additional primers were also used in the development of the *Pam*^cKO/cKO^ line, to verify the cloning strategy at each step of the process. Wildtype mice yielded a 194 nt product with primers A and B (A:B), and no product when primer A was paired with C (A:C) (**Suppl.Fig.1A**). When the floxed allele was present, addition of the loxP site yielded an A:B product of 248 nt, but no A:C product. In a full knockout tissue (such as atrium from *Pam*^Myh6-cKO/cKO^), A:B yielded no product, but A:C yielded a 310 nt product. A *Pam*^Myh6-cKO/+^ mouse yielded two bands (194 and 310 nt) with the A:B:C mixture in Cre-expressing tissues. Designing a three-primer screen for the Emx1-Cre mice first required sequencing the genomic segment extending from the end of the Emx1 gene into Cre-recombinase, and then creating a reverse primer unique to the Emx1-Cre allele. The three-primer screen then yielded a 160 nt product from the Emx1 wildtype allele and a 230 nt product from Emx1-Cre (**Suppl.Fig.1B**). As per the JAX website, Emx1-Cre mice were maintained as homozygous breeding pairs, but Myh6-Cre mice were kept as heterozygotes. Similarly, genotyping the Myh6-Cre mice was performed with a three primer set, with a common forward primer and two reverse primers, one from the wildtype at 214 nt and one from the Myh6-Cre at 300 nt (**Suppl.Fig.1C**).

**Supplemental Results**

**Immunocytochemistry controls**

Controls for immunostaining included antibody specificity on Western blot analyses and ablation of signal in knockout tissue (**Fig.2C,D,E**). In addition, signal that remained after inclusion of antigenic peptide (Exon 16) along with PAM antibody JH629 (**Suppl.Fig.4C, 5C, 6C, 7C**) or remained when non-immune rabbit IgG replaced primary antibody was taken as background (**Suppl. Fig.4B, 5B, 6B, 7B**). Use of these controls identified PAM immunostaining that was specific.

**Further RNAseq analyses**

Ingenuity Pathway Analysis yielded a number of interesting networks centered on transcription factors whose expression levels were unchanged but which are known to be regulated by phosphorylation (**Supp.Fig.8**). The Crem pathway includes two of the transcripts from the circadian group which increased with the loss of *Pam*, Bhlhe40 and Per1, along with Ciart. The Stat6 pathway included several strongly regulated transcripts: Ano1 (Anoctamin 1 or Tmem16a, a calcium activated chloride channel), Arntl2 (Aryl hydrocarbon receptor nuclear translocator-like 2), Gadd45g (Growth arrest and DNA-damage-inducible 45 gamma), Il18r1 (Interleukin 18 receptor 1) and Usp2 (Ubiquitin specific peptidase 2). The Epas1 pathway includes Apln and Bhlhe40 from the pathways in **Fig.8**, as well as the Ca^++^ ion channel Cacna1a.

**Supplemental Figures**

**Supplemental Fig.1.** **Design of *Pam*^Emx1-cKO/cKO^ mice and *Pam*^Myh6-cKO/cKO^ mice**. **A.** The region on mouse chromosome 1 around *Pam* Exons 2-4 is depicted (not to scale) (1). The positions of the loxP sites are depicted as blue triangles; the G418 resistance sequence was removed by Cyagen to create the F1 generation of mice, with the one remaining Rox site shown as a brown diamond. PCR primers A, B and C (**Suppl.Table 1**) are depicted in green, light blue and red, respectively. For simplicity, the gel at the right shows the results of genotyping with pairs of primers; all three primers were normally present in the reaction tube. **B.** The structure of the genomic region encompassing the *Emx1* stop codon and proximal 3’-untranslated region on chromosome 6 is depicted at the top. The sequenced *Emx1-IRES-Cre* genomic region is diagrammed at the bottom (2). The green and light blue primers yielded a 160 nt product from the endogenous *Emx1* gene, while the green and red primers yielded a 230 nt product only from the *Emx1-IRES-Cre* sequence. An earclip screening gel is shown at the right. **C.** The structure of the *Myh7-Myh6* genomic boundary region on chromosome 14 is depicted, along with the *Myh6-Cre* transgene (3). The green and light blue primers yielded a 214 nt product only from the endogenous mouse *Myh6* gene, while the green and red primers yielded a 300 nt product only from the *Myh6-Cre* transgene, as seen in the gel on the right.

**Supplemental Fig.2.** **Further analyses of the tissue-specific knockout mice**. **A.** Pups born and weaned from mating *Pam*^Emx1-cKO/+^ mice with *Pam*^Emx1-cKO/+^ mice were found in the expected Mendelian ratios (N=226; N.S. by 2-way ANOVA). **B.** Similarly, mice were born and weaned in numbers not significantly different from those predicted based on Mendelian inheritance from pairing *Pam*^Myh6-cKO/+^ mice with *Pam*^cKO/+^ mice (N=198; N.S. by 2-way ANOVA). **C.** Quantitative polymerase chain reaction (qPCR) was performed on atrial RNA from WT and *Pam*^Myh6-cKO/cKO^ mice using primers designed to verify the absence of transcripts containing Exons 2 and 3 in *Pam*^Myh6-cKO/cKO^ samples. **D.** qPCR using primer pairs specific for the major splice variants of mouse PAM (PAM1, PAM2 and PAM3) was used to determine the major isoforms expressed in the adult atrium. Data for *Pam* were normalized to levels of *Gapdh* in the same samples. Similar results were obtained in repeat assays with independent samples.

**Supplemental Fig.3.** **Controls for immunofluorescence studies localizing sites of PAM expression in TdTomato reporter mice - CA1. A.** PAM-positive cells (green) in the CA1 region of the hippocampus; PAM antibody JH629 was used. **B.** IgG control staining in the same region. **C.** Peptide (PAM exon 16) blocking control in the same region. Based on TdTomato fluorescence, the most intensely stained PAM-positive neurons (marked with a yellow arrow) were not Emx1-positive (TdTomato-expressing) excitatory neurons.

**Supplemental Fig.4. Controls for immunofluorescence studies localizing sites of PAM expression in TdTomato reporter mice - CA3.** Experimental design was as described for **Suppl. Fig.3**. **A.** PAM-positive cells in the CA3 region. **B.** IgG control. **C.** Peptide (PAM exon 16) blocking control. Most of the PAM staining seen in Panel A was in mossy fiber nerve endings.

**Supplemental Fig.5. Controls for immunofluorescence studies localizing sites of PAM expression in TdTomato reporter mice - Cortex layer 2.** Experimental design was as described for **Suppl. Fig.3**. **A.** PAM-positive cells in Layer 2. **B.** IgG control. **C.** Peptide (PAM exon 16) blocking control. Yellow arrows mark PAM-positive neurons which were TdTomato-positive; white arrow marks PAM-positive neuron which was not TdTomato-positive.

**Supplemental Fig.6. Controls for immunofluorescence studies localizing sites of PAM expression in TdTomato reporter mice - Cortex layer 5.** Experimental design was as described for **Suppl. Fig.3**. **A.** PAM-positive cells in Layer 5. **B.** IgG control. **C.** Peptide (PAM exon 16) blocking control. PAM-positive neurons marked with yellow arrows were not TdTomato-positive.

**Supplemental Fig.7. TdTomato is expressed in Myh6-expressing atrial myocytes**. Adult mice expressing a TdTomato Cre-reporter construct and Myh6-Cre were perfusion fixed; nuclei in sections of the left atrium were visualized using Hoechst stain (blue). Epifluorescence images of TdTomato fluorescence and Hoechst stain were taken using a Nikon TE300 microscope with a 10X-objective. Essentially all of the atrial myocytes expressed TdTomato.

**Supplemental Fig.8. Additional significantly altered networks in WT vs. *Pam*^Myh-cKO/cKO^ atria. A. Crem.** *Crem* (cAMP responsive element modulator) levels were unaltered (FPKM=25). Network includes 9 transcripts increased in *Pam*^Myh6-cKO/cKO^ atria and decreased in pituitary cells with elevated PAM (4) [*], 1 transcript not expressed in pituitary [0], and only 1 transcript in disagreement with the pituitary result [X] and also in disagreement with the Crem network prediction (yellow arrow). **B. Stat6.** *Stat6* (signal transducer and activator of transcription 6) levels were unaltered (FPKM=70). Network includes 4 transcripts regulated similarly in pituitary cells, 1 transcript unchanged in pituitary [N], 2 not expressed in pituitary and only 2 in disagreement between atrium and pituitary. **C. Epas1.** *Epas* (Epithelial PAS domain protein 1, also known as *Hif2α*) levels were unaltered (FPKM=156). Includes 3 transcripts regulated as in pituitary, 3 not expressed in pituitary (including *Apln*), and 3 in disagreement with the pituitary result. Code: *, significant and in agreement in pituitary cells (4); N, no change in pituitary cells; 0, not expressed above FPKM=2 in pituitary cells; X, opposite response in pituitary cells. Regulated transcripts are described in more detail in **Suppl.Table 4**.

**Supplemental Table 1. Primers**

| Genotyping primers | | | |
| --- | --- | --- | --- |
| target | forward | reverse | Size (nt) |
| PAM 5’ loxP | ATCGTGGTCTCTAAGTGCTTTGTC | GGTTGTTCACCCTGCCTTCCTTTTC | WT 192  LoxP 248 |
| PAM 3’ loxP |  | TGTTTTTAATGACATACATAAATCCCCGCACTG | Deletion 310 |
| Myh6-no Cre | ATGACAGACAGATCCCTCCTATCTCC | TTCCCGGGACACGACCTTGGC | WT 214 |
| Myh6- Cre |  | GCGAACCTCATCACTCGTTGCATC | Cre 300 |
| Emx1-no Cre | GGGGAGGACATTGATGTCACCTCC | CTGTGGGGCCGTGACTCGAG | WT 160 |
| Emx1-Cre |  | CGGAATTCCGGTCTCCCTATAGTGAG | Cre 230 |
| qPCR Primers | | | |
| Anp | CAAGAACCTGCTAGACCACCTGGAG | CCTCGGGGAGGGAGCTAAGTG | 120 |
| Bnp | TCCTAGCCAGTCTCCAGAGCAATTCAAG | GGGTGTTCTTTTGTGAGGCCTTGGTC | 120 |
| Pam ex2 | CCCGCTGCCCCGGTCCTGCGCGGA | CTCTTAAAGACAGAAAGTGGGCTTCTGAAG | 120 |
| Pam ex3 | GTTTAAAGAAACTACCAGATCATTTTCCAA | CTCTTTAGGTGTGACCCCAGGCATG | 120 |
| Pam1 | GAGGAAGCCTTCGAGCAGGGTG | CGTTTGCAATCTCAGCTACCAGGTCTG | 120 |
| Pam2 | AGGAGGAAGCCTTCGAGCAGGATTTC | AAATCACAAGGTTATTCTTAGAATCCAGGGCC | 120 |
| Pam3 | TGGCATTGAGGTCCCGGAAATCAAAGATC | CTTGCAAAGAAATTTCCTAGATTTAAGCCGCTG | 120 |

**Supplemental Table 2**. **Cell counts.** PAM-positive neurons in cortical layers 2 and 5 in TdTomato mice and in *Pam*^Emx1-cKO/cKO^ mice were counted and compared to total cell number, as determined by Hoechst stain). GAD-positive neurons were also counted in *Pam*^Emx1-cKO/cKO^ mice.

| **Cortical Layer** | **TdTomato**  **+ PAM** | **TdTomato**  **no PAM** | **PAM only** | **GAD only** | **GAD + PAM** | **Total PAM** |
| --- | --- | --- | --- | --- | --- | --- |
| **2 (TdTomato)** | **56.0 ± 9.5** | **0** | **11.3 ± 7.7** | **-** | **-** | **67.3 ± 1.8** |
| **2 (WT)** | **-** | **-** | **63.2** | **0** | **2.7** | **66.0** |
| **2 (Emx1-Cre)** | **-** | **-** | **7.8** | **0** | **1.9** | **9.7** |
| **5 (TdTomato)** | **44.3 ± 6.4** | **0** | **9.1 ± 5.4** | **-** | **-** | **53.4 ± 1.1** |
| **5 (WT)** | **-** | **-** | **36.9** | **0** | **9.2** | **46.1** |
| **5 (Emx1-Cre)** | **-** | **-** | **4.1** | **0** | **3.5** | **7.6** |

A “-“ marks measurements that do not apply to the mice under study. For the TdTomato images, two mice were perfused, sectioned and imaged; the images had extremely similar percentages of cells (Hoechst) which were PAM-positive, while the proportion of PAM-positive neurons which expressed TdTomato was more variable. The error shown is the range.

**Supplemental Table 3. FPKM data for atrial samples from adult WT and *Pam*^Myh6-cKO/cKO^ mice.** RNA samples from 25 adult mice [12 WT (7 male, 5 female) and 13 *Pam*^Myh6-cKO/cKO^ (5 male and 8 female)] were sequenced and subjected to DESeq2 differential expression analysis. FPKM values are shown for the WT males (WM1-WM7), WT females (WF8-WF12), *Pam*^Myh6-cKO/cKO^ males (KM13, KM01, KM03, KM05, KM07), and *Pam*^Myh6-cKO/cKO^ (females (KF14-KF21), along with the DESeq2 p-value, average FPKM for WT and *Pam*^Myh6-cKO/cKO^ samples, and Log2(WT/*Pam*^Myh6-cKO/cKO^).

**Supplemental Table 4**. **Transcript network members.**

All the transcripts displayed in **Fig.8** and **Suppl.Fig.8** are listed alphabetically. Red type means decreased in *Pam*^Myh6-cKO/cKO^ mice, while blue type means increased in *Pam*^Myh6-cKO/cKO^ mice (same color code as in **Fig.8**). An ‘X’ in the name box means that the changes in transcript level in atrium (this work) and pituitary (4) do not agree.
