## Supplementary figures and images for "Identifying Novel Roles for Peptidergic Signaling in Mice"

### Supplemental Figures

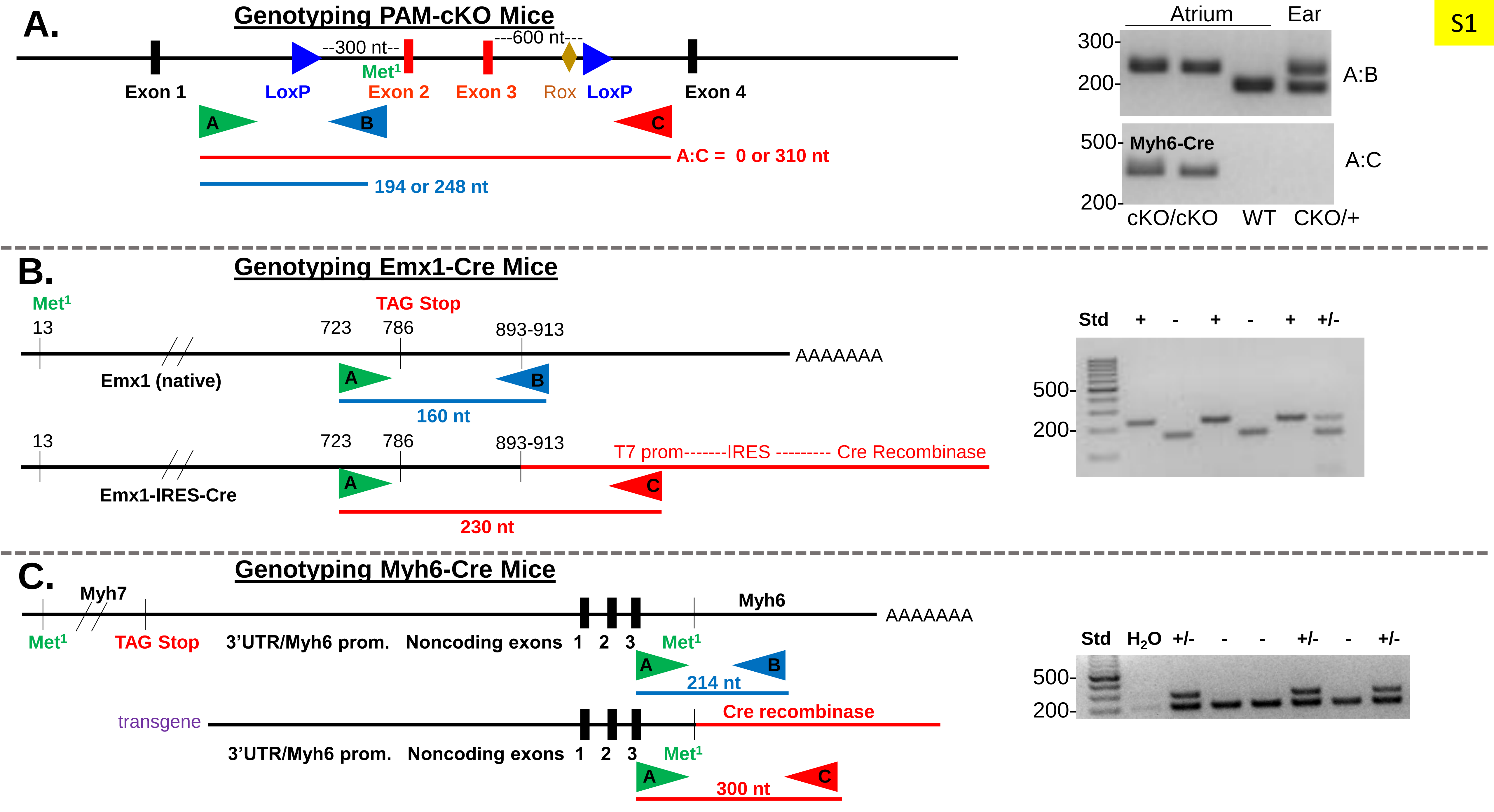

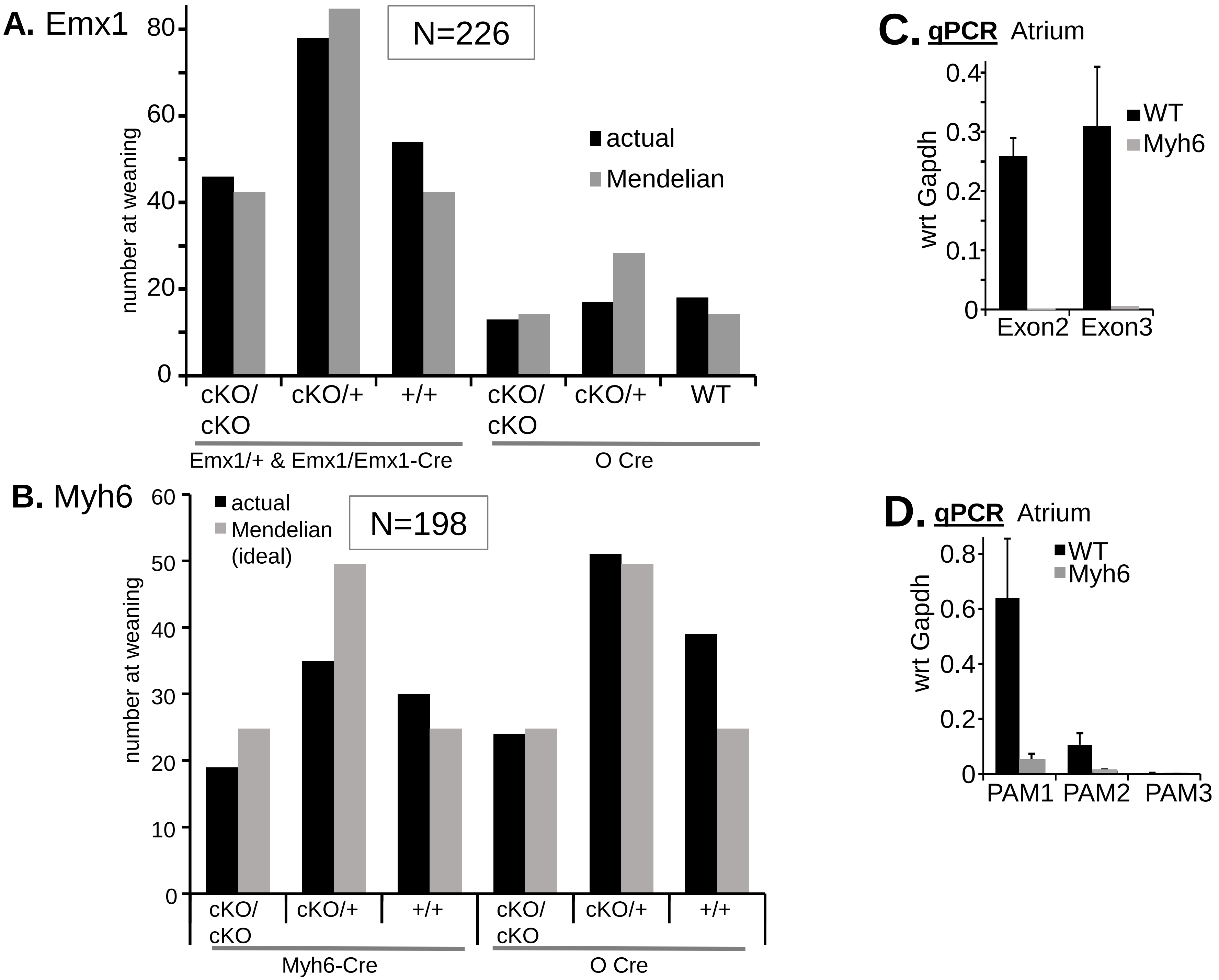

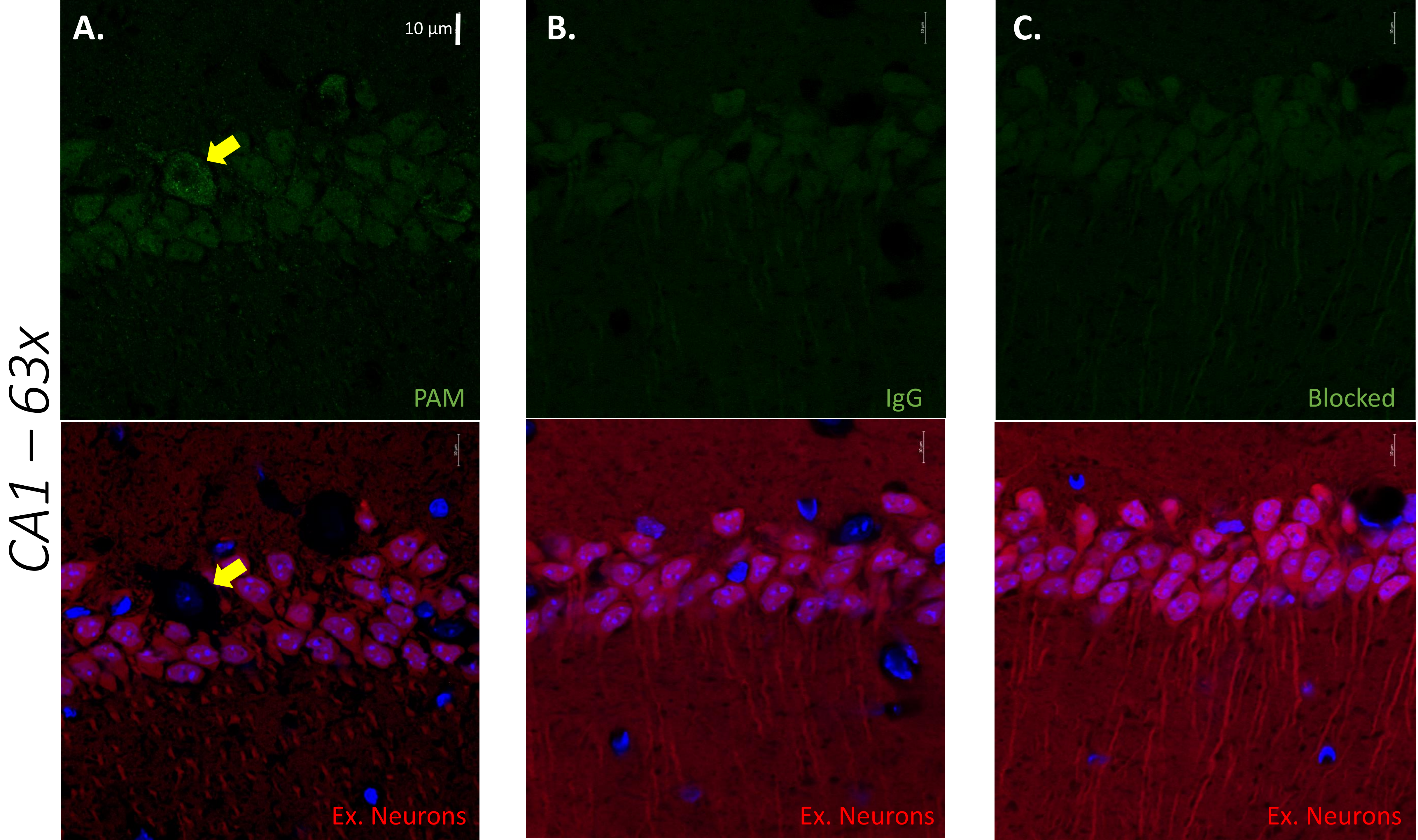

S4

CA3 – 63x

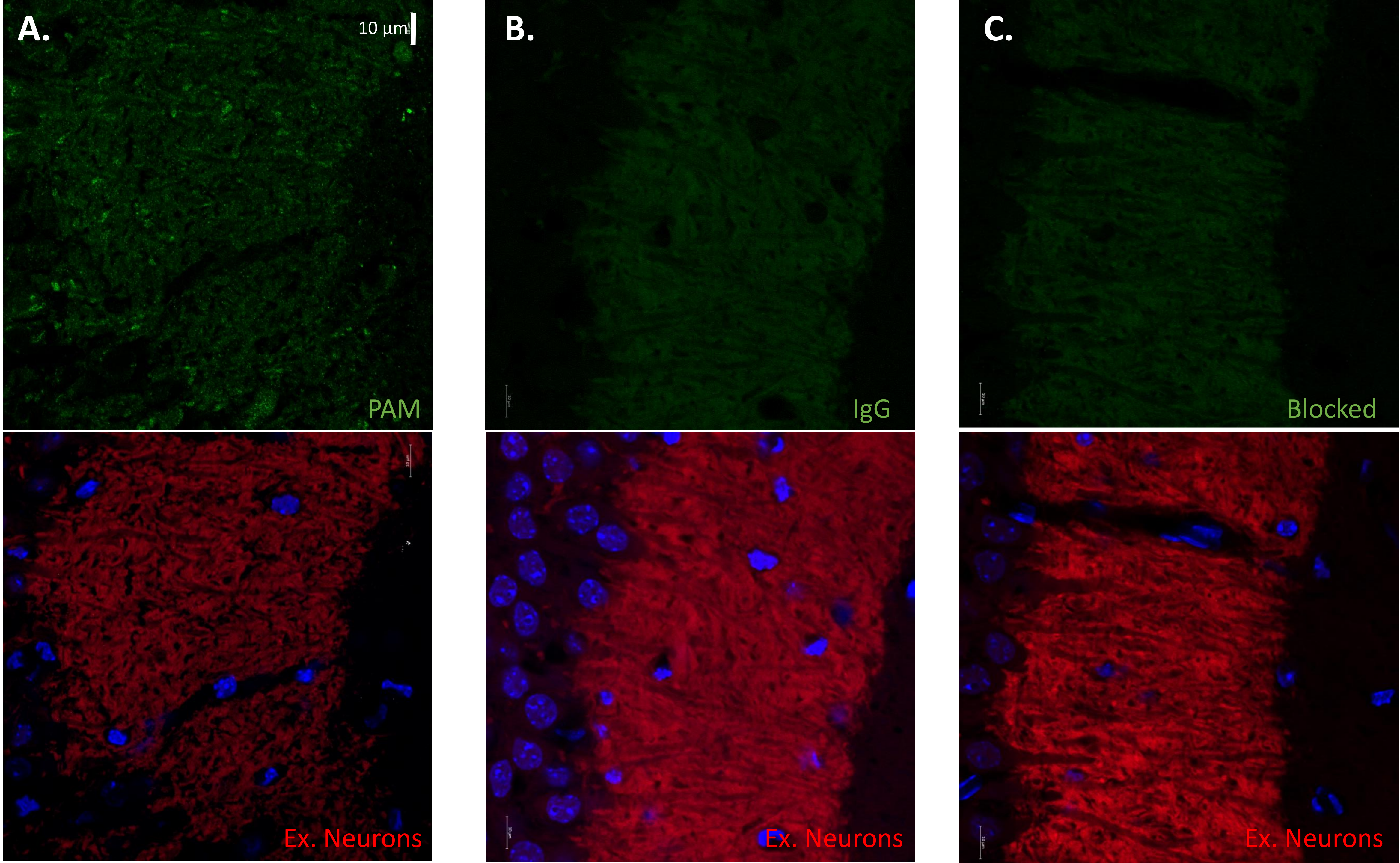

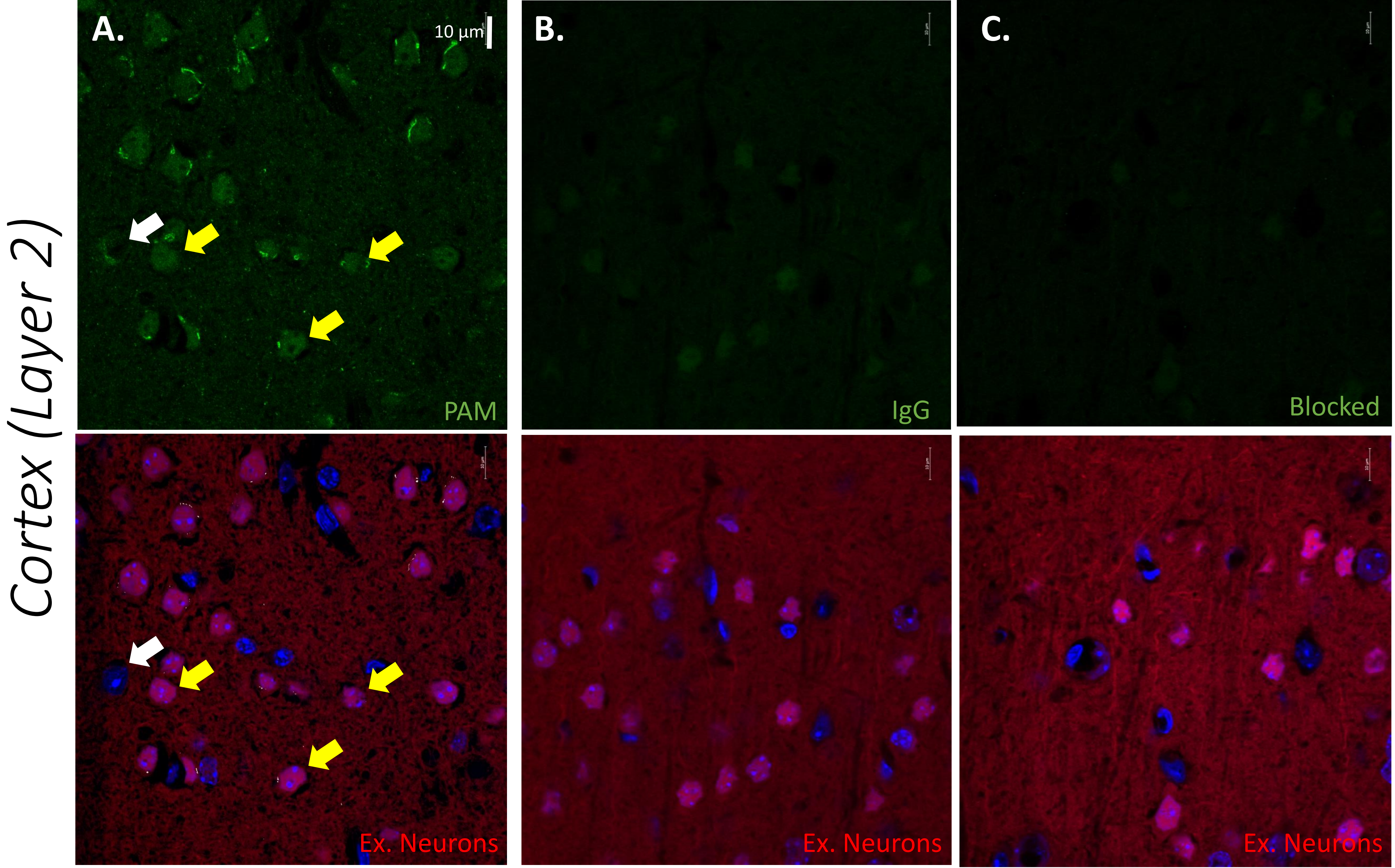

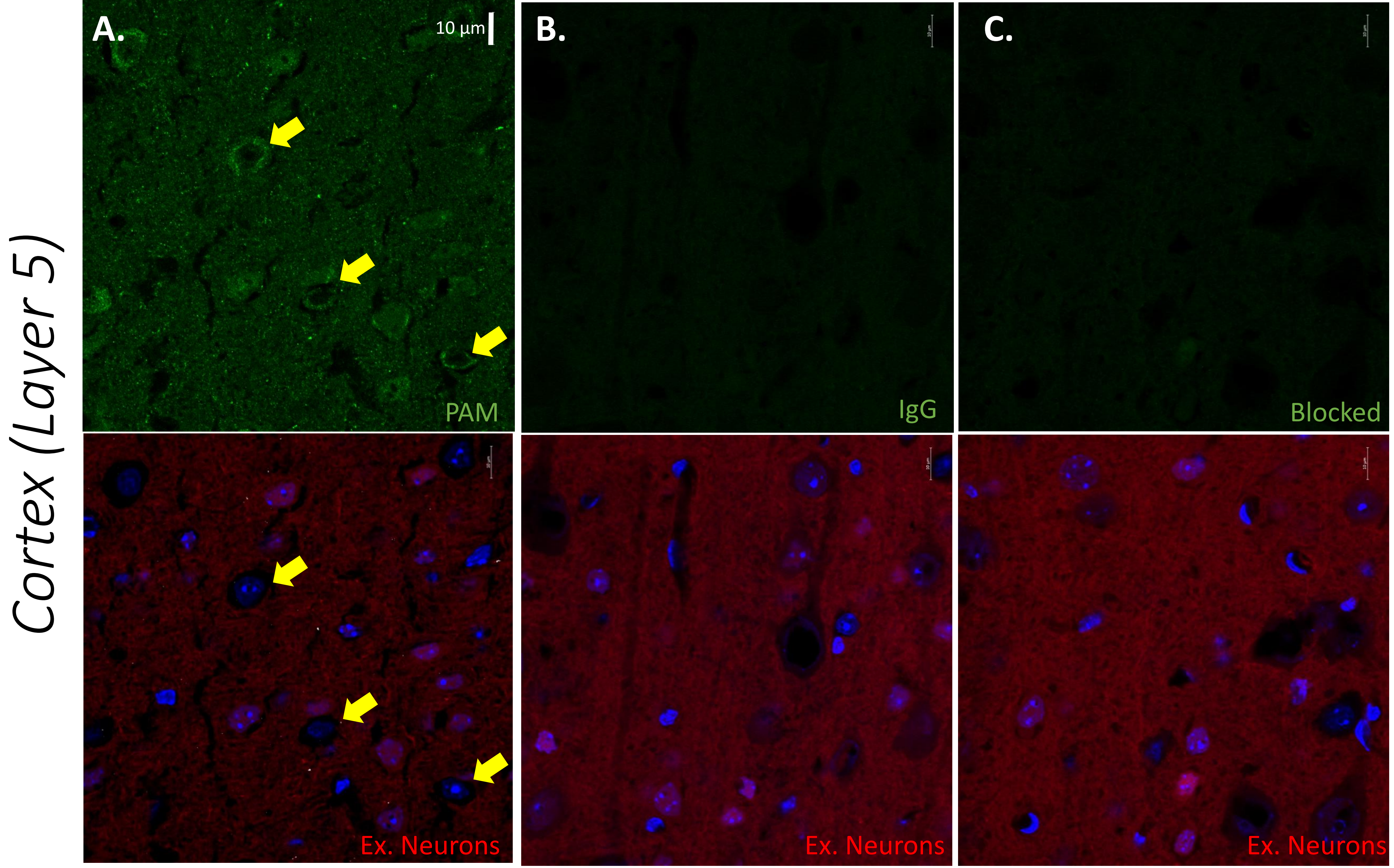

S7

#### *Atrium*

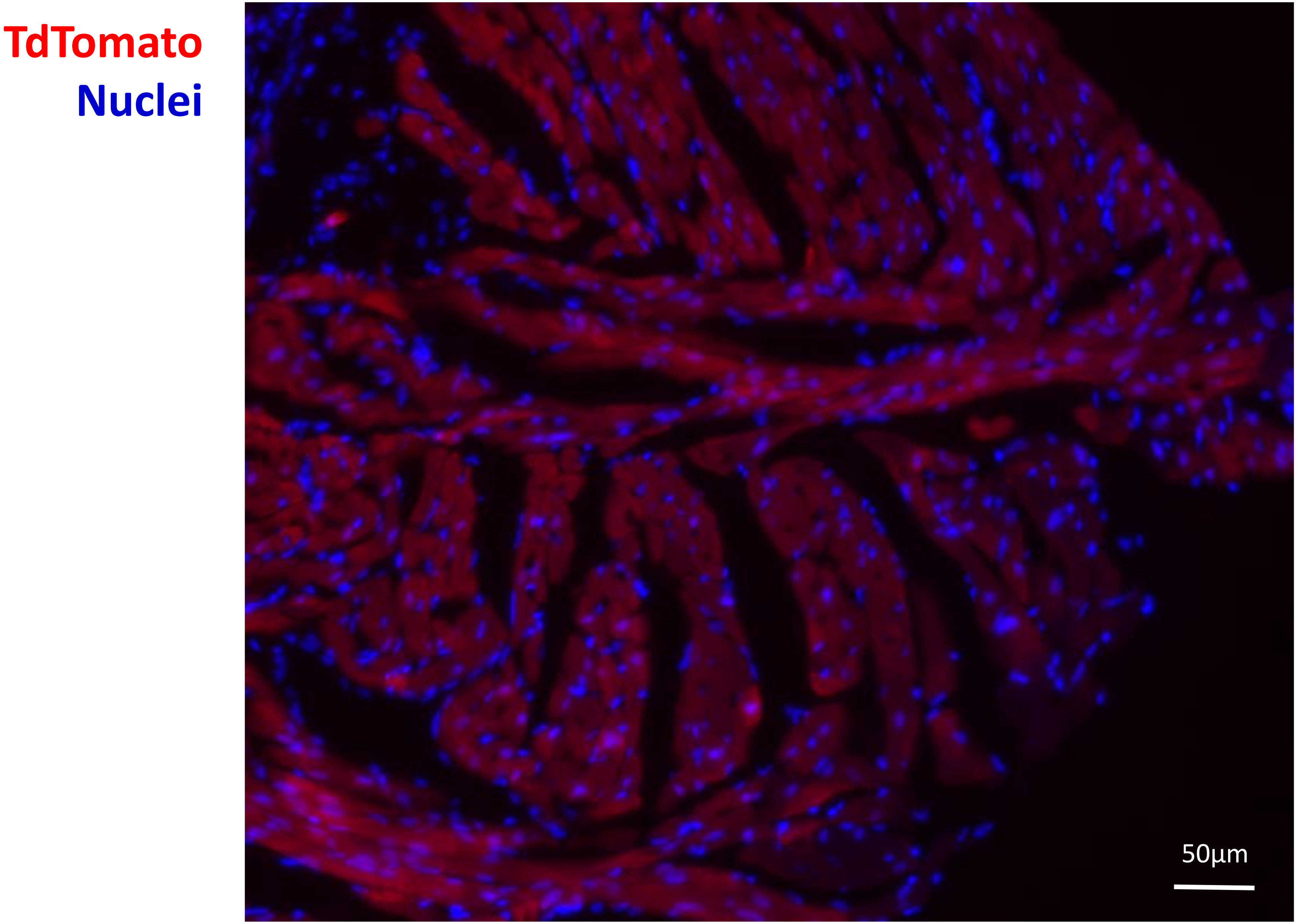

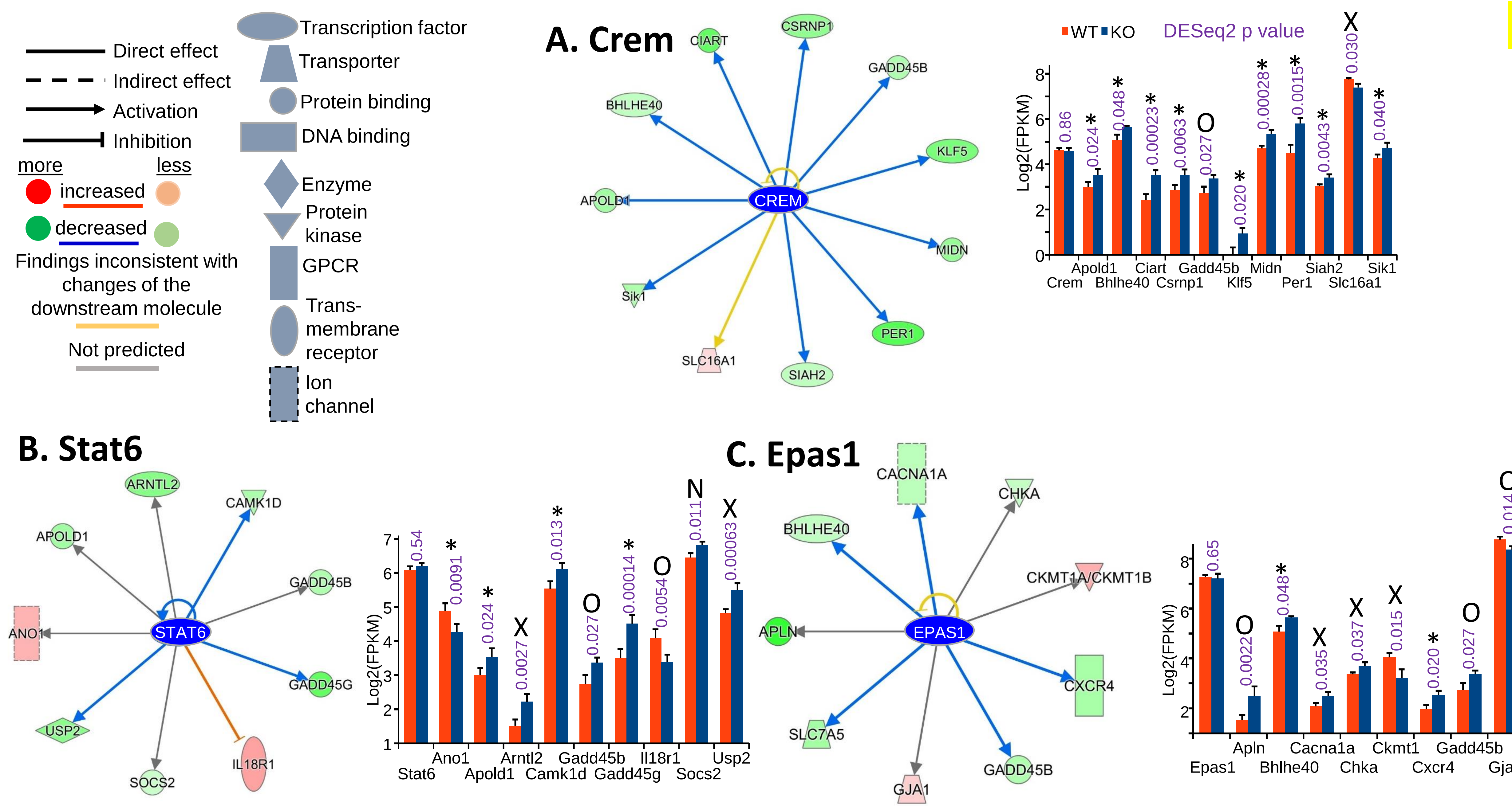
