## Supplemental Table 4 for "Identifying Novel Roles for Peptidergic Signaling in Mice"

**Transcript changes**	color code: down in PAMKO, up in PAMKO

| **transcript** | **Most have same direction CHANGE with PAM in atrium & pituitary; X = Opposite direction** |
| --- | --- |
| Atf4 | Activating transcription factor 4. A major regulator of amino acid metabolism, induced during starvation; can be pro=apoptotic. Heart RPKM 27; atrium FPKM 110. |
| Ano1 | Anoctamin 1, calcium activated chloride channel (Tmem16a). Modulates arterial contractility; bile acids activate Cl- secretion via Tmem16a; stained in lots of presynaptic endings. Heart RPKM 3; atrium FPKM 33. |
| Apln | Apelin. Mice lacking Apln develop heart failure, ligand for G-protein coupled receptor Aplnr (also called Apj). Heart RPKM 3; atrium FPKM 3. (Aplnr FPKM 14) |
| Apold1 | Apolipoprotein L domain containing 1 Verge). Verge= vascular early response gene crucial to developmental angiogenesis, and adult cerebral ischemia. Heart RPKM 1.6; atrium FPKM 9. |
| Arntl  X | Aryl hydrocarbon receptor nuclear translocator-like (Bmal1). Forms heterodimer with Clock; activates Per 1,2,3 and Cryptochromes Cry1,2. Then feedback by Per and Cry represses via negative feedback. Heart RPKM 1.6; atrium FPKM 21. |
| Arntl2  X | Aryl hydrocarbon receptor nuclear translocator-like-2 (Bmal2). Forms heterodimer with Clock and Hif1a hypoxia factor. Heart RPKM 0.24; atrium FPKM 3. |
| Bhlhe40 | Basic helix-loop-helix family member e40 (Dec1). Interacts with Arntl, competes for E-box on Per1 promoter, suppresses Clock/Arntl activation of Per1. Heart RPKM 9; atrium FPKM 38. |
| Cacna1a  X | Calcium channel, voltage dependent, P/Q type, Alpha 1a subunit (Cav2.1). Role mostly defined in nervous system (presynaptic vesicle release). Heart RPKM 0.4; atrium FPKM 4. |
| Camk1d | Calcium/calmodulin-dependent protein kinase 1d (Cklik). Translocates to nucleus when intracellular Ca++ influx increases. Heart RPKM 0.75; atrium FPKM 51. |
| Chka  X | Choline kinase alpha. Regulated by circadian clock; makes phosphatidylcholine, not rate-limiting in adult; Heart RPKM 0.7; atrium FPKM 10. |
| Ciart | Circadian associated repressor of transcription (Gm129; Chrono). Modulates circadian gene expression; a core component of circadian clock; functions in preimplantation embryo. Heart RPKM 1.3; atrium FPKM 6. |
| Ckmt1  X | Creatine kinase, mitochondrial 1, ubiquitous. Chr 2; CNS KO 🡪 problems in thermoregulation and learning. Heart RPKM 0; atrium FPKM 18. |
| Clock | Circadian locomotor output cycles kaput. Transcription factor, central to circadian rhythms, forms dimer with Arntl. Involved in behavioral changes, obesity. Heart RPKM 4; atrium FPKM 177 |
| Csrnp1 | Cysteine serine rich nuclear protein 1. Induced by Axin; Wnt pathway repressed by Axin; Csrnp1 is a tumor suppressor; multiple transcripts. Heart RPKM 3; atrium FPKM 8. |
| Cxcr4 | Chemokine (C-X-C motif) receptor 4. Functions in hippocampal development, bone resorption. Heart RPKM 7; atrium FPKM 4. |
| Gadd45b | Growth arrest and DNA-damage-inducible 45 beta. Heart RPKM 3; atrium FPKM 8 |
| Gadd45g | Growth arrest and DNA-damage-inducible 45 gamma (Ddit2). Pax5 upregulates; accumulates during myocardial infarction; a cold-inducible activator of Ucp1 uncoupling protein 1; activates oxidation in brown fat.. Heart RPKM 7; atrium FPKM 13. |
| Gja1 | Gap junction protein alpha 1 (Cx43). Involved in gap junction responses to hypoxia and low glucose. Heart RPKM 61; atrium FPKM 450. |
| Gnas | Guanine nucleotide binding protein, alpha stimulating. Many splice forms & different protein isoforms, 5’ methylation affects expression, has an antisense transcript. In most tissues; heart RPKM 57; atrium FPKM 5. |
| Gpx3 | Glutathione peroxidase 3. Catalyzes reduction of hydroperoxides including H_2_O_2_ by GSH; this isozyme is a secreted selenoprotein. Heart RPKM 135, Kidney 12,000; atrium FPKM 1930. |
| Hdac4 | Histone deacetylase 4. Heart overexpression causes myocardial ischemia. In most tissues; heart RPKM 2.6; atrium FPKM 16. |
| Il18r1 | Interleukin 18 receptor 1. Much studied in inflammation and neuronal development. Heart RPKM 0.1; atrium FPKM 20 |
| Klf5 | Kruppel-like factor 5. Suppresses Erk signaling; essential for mammary gland development; essential in lung, liver, colon development. Heart RPKM 0; atrium FPKM 1. |
| Midn | Midnolin. Has an ubiquitin-like domain. High expression in embryonic brain. Alternate mRNA splicing, multiple protein isoforms. Heart RPKM 21; atrium FPKM 27. |
| Per1 | Periodic circadian clock 1. Circadian in suprachiasmatic nucleus. Upregulated by Clock/Arntl heterodimers, then represses its own upregulation in Per/Cry heterodimers suppressing Clock/Arntl. Heart RPKM 6; atrium FPKM 30. |
| Per2 | Periodic circadian clock 2. Same story as Per1. Heart RPKM 2; atrium FPKM 20. |
| Per3 | Periodic circadian clock 3. Same story as Per1 and Per2. Heart RPKM 3; atrium FPKM 34. |
| Prkab2 | Protein kinase AMP-activated non-catalytic subunit beta-2. Heart RPKM 10; atrium FPKM 46. |
| Prkag2  X | Protein kinase, AMP-activated, gamma 2 noncatalytic subunit. Activation decreases sinoatrial pacemakers, lowers heart rate, lowers ryanodine-driven Ca++ release from intracellular stores. Heart RPKM 4; atrium FPKM 105. |
| Prkar2b  X | Protein kinase, cAMP dependent regulatory, type 2 beta. Deficiency increases expression of Ucp1, other thermogenic genes, induces brown-like fat in white adipose; disruption increases resting metabolic rate. Heart RPKM 2.4; atrium FPKM 20. |
| Rras2  X | Ras related 2 (Tc21). Fatty-acylated protein, regulates Tgf-beta signaling. Heart RPKM 5; atrium FPKM 70. |
| Siah2 | Siah E3 ubiquitin protein ligase 2 (Sinh2). Important for early adipogenic pathway; obesity-induced tissue inflammation. Heart RPKM 3; atrium FPKM 8. |
| Sik1 | Salt inducible kinase 1. Induced by alcohol; KO of Sik1 causes increased blood pressure; regulates intercellular junction stability; senses intracellular sodium. Heart RPKM 6; atrium FPKM 20. |
| Slc16a1  X | Solute carrier family 16 (monocarboxylic acid transporters), member 1 (Mct1). Heart RPKM 45; atrium FPKM 220. |
| Slc7a5 | Solute carrier family 7 (cationic amino acid transporter, y+ system) member 5 (Lat1). Regulates Kv1.2 channels, relevant to epilepsy; stretching stimulates Slc7a5 expression and subsequent amino acid uptake. Heart RPKM 2; atrium FPKM 3. |
| Socs2 | Suppressor of cytokine signaling 2. Roles in immune function, nervous system; in heart, lack of Socs2 blocks responses to increased Ca^++^. Heart RPKM 2; atrium FPKM 90 |
| Tgfb1 | Transforming growth factor beta 1. Secreted; forms heterodimers with other Tgf-beta family. Heart RPKM 7; atrium FPKM 29 |
| Usp2  X | Ubiquitin specific peptidase 2 (Ubp41). Clock output effector, regulates body Ca++; represses MC receptor and thus suppresses aldosterone responses; a central regulator of metabolic rate; not directly involved in Na balance or blood pressure; rhythmic expression circadian. Heart RPKM 6.5; atrium FPKM 29. |

Heart RPKM from NCBI website; atrium FPKM from Suppl.Table 3 (WT value)
